# High-resolution mapping of cancer cell networks using co-functional interactions

**DOI:** 10.1101/369751

**Authors:** Evan A. Boyle, Jonathan K. Pritchard, William J. Greenleaf

## Abstract

Powerful new technologies for perturbing genetic elements have expanded the study of genetic interactions in model systems ranging from yeast to human cell lines. However, technical artifacts can confound signal across genetic screens and limit the immense potential of parallel screening approaches. To address this problem, we devised a novel PCA-based method for eliminating these artifacts and bolstering sensitivity and specificity for detection of genetic interactions. Applying this strategy to a set of >300 whole genome CRISPR screens, we report ~1 million pairs of correlated “co-functional” genes that provide finer-scale information about cell compartments, biological pathways, and protein complexes than traditional gene sets. Lastly, we employed a gene community detection approach to implicate core genes for cancer growth and compress signal from functionally related genes in the same community into a single score. This work establishes new algorithms for probing cancer cell networks and motivates the acquisition of further CRISPR screen data across diverse genotypes and cell types to further resolve the complexity of cell signaling processes.

## Introduction

Understanding the complex biological underpinnings of human disease has long been a goal of network biologists (Barabási & Oltvai, 2004; Barabási *et al*, 2011). Because genes vary in their role and importance across diverse cell types, it has become increasingly clear that characterizing tissue- and cell type-specific regulation of chromatin accessibility (Roadmap Epigenomics Consortium *et al*, 2015;Breeze *et al*, 2016), chromosome looping (Javierre *et al*, 2016; Mumbach *et al*, 2017), and gene expression (GTEx Consortium *et al*, 2017) will be central to developing a coherent understanding of disease etiology. The differences in biological pathway importance across tissues is especially vexing when modeling diseases such as cancer that specifically exploit tissue-specific pathways and preferentially acquire mutations to regulate them. Set against these challenges, the advent of new genetic perturbation systems scalable to the size of the human genome offers an unprecedented opportunity for the study of cancer cell networks and associated tissue-specific signaling paradigms that do not exist in single-celled model organisms like yeast (Gilmore, 2006; Fontana *et al*, 2010).

Genome-wide CRISPR screens (Shalem *et al*, 2014; Wang *et al*, 2014) have already enabled insights into cell trafficking (Gilbert *et al*, 2014), drug mechanism of action (Shalem *et al*, 2014; Wang *et al*, 2014; Doench *et al*, 2016), and infectious disease (Gavory *et al*, 2018; Park *et al*, 2017). Yet, while these methods allow every gene to be perturbed and scored for its effect on a phenotype of interest, this score does not provide direct insight into the logic of the biological pathways involved in mediating the phenotype. Instead, these screens report one dimensional vectors of values: each gene falls on a single spectrum from dis-enriched to enriched. Determining the cellular logic that integrates effects across genes requires either specialized experimental design or extensive post-processing of high-throughput screen data. Presently, there are three prominent strategies that assign pathways to genes and explain interactions between genes: computing enrichment of hits in curated gene sets, quantifying effect sizes of combinations of genetic perturbations, and analyzing diverse CRISPR screens performed in parallel to identify genes with correlated effect sizes. Enrichment in curated gene sets can clarify the biological processes involved by implicating cell pathways or compartments, but interpreting enrichments can be extremely difficult (Rhee *et al*, 2008; Timmons *et al*, 2015). Thus, we will focus on the two experimentally-driven alternatives.

Combinatorial sgRNA platforms (Bassik *et al*, 2013; Kampmann *et al*, 2015) disrupt multiple genes in the same cells to identify epistatic interactions that can unmask pathway logic. These platforms are the successors of double knockout array technology that has broadened the set of yeast genes thought to participate in genetic interactions to 90% of all genes (Tong *et al*, 2004; Costanzo *et al*, 2010; 2016). While these methods have the power to directly test hypothesized genetic interactions, technological constraints have limited individual combinatorial sgRNA studies to measuring interactions for only a small fraction of all pairs of human genes (Ogasawara *et al*, 2015; Han *et al*, 2017; Shen *et al*, 2017; Wong *et al*, 2016). Ascertaining appropriate gene pairs for such phenotyping is not trivial, especially for synthetic interactions where neither genetic perturbation exhibits an effect on its own, although algorithms to overcome this challenge are under development (Medina & Goodin, 2008; Deshpande *et al*). Interpretation of measured interactions is also complicated by the fact that the extent to which genetic interactions persist across cell types or samples is unknown.

Parallel screening designs approach the identification of interacting genes in a fundamentally different manner. All genes of interest, potentially comprising the entire genome, are screened in a diverse panel of cell lines, and the perturbation effect sizes across these cell lines are recorded as a gene perturbation profile for every gene (Figure 1a, b). The distinct genetic and epigenetic features of each cell line modify its susceptibility to disruption of pathways, organelles, or even individual protein complexes. In general, two genes that have correlated gene perturbation scores across many cell lines are inferred to be functionally related, with greater correlation implying greater shared function. Data from only 6 cell lines sufficed to verify essential gene pathways in one study (Hart *et al*, 2015). A larger panel of 14 parallel AML line CRISPR screens allowed more systematic validation of cancer metabolic and signaling pathways (Wang *et al*, 2017). More recently, the availability of 342 published CRISPR screens in cell lines drawn from diverse cell lineage and mutational backgrounds has invited even broader surveys of co-essentiality (Meyers *et al*, 2017). One study jointly examining RNAi screen data dissects the structure of essential protein complexes (Pan *et al*, 2018), and a second, currently available on a preprint server, independently examines the organization of cancer growth pathways (Kim *et al*, 2018). In all these cases, genetic interactions identified from correlated gene profiles operated on multiple levels of cellular regulation, validating parallel screening as a powerful tool for reconstructing cell networks.

**Figure 1:**
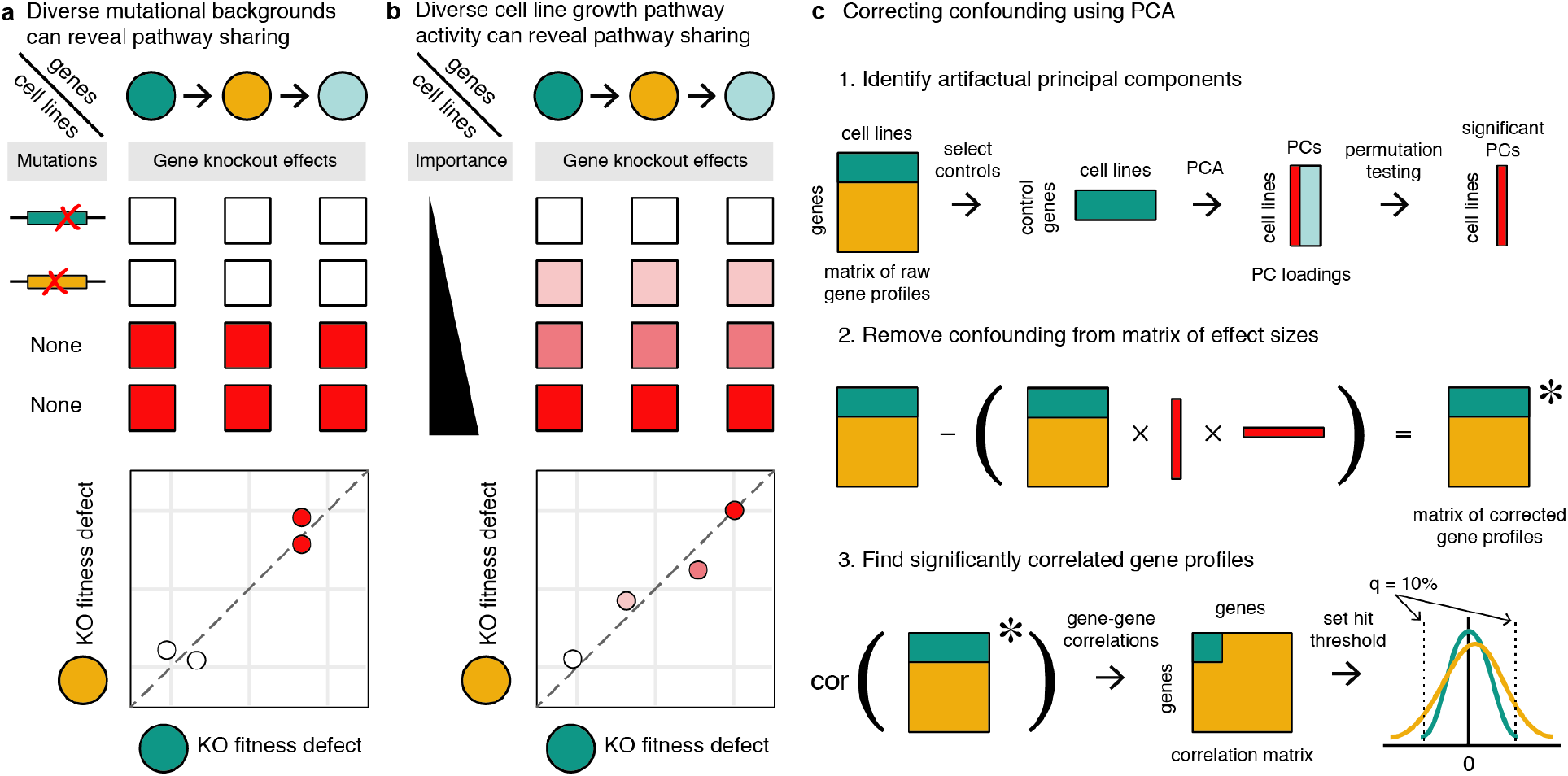
Detecting co-functional interactions from parallel genetic perturbation screens. Diversity in **a)** mutational background and **b)** pathway activity produce correlated gene essentiality profiles of knockout effect sizes for members of the same pathway. Cell lines with inactivating mutations in or downregulation of biological pathways are impervious to gene knockout. Discrete and continuous differences across cell lines can thus be summarized by a correlation coefficient. **c**) Nonspecific sources of variation across cell lines can be removed by learning principal components from a set of control genes that should not contain biological signal and subtracting these components from the raw gene profiles.

While effective, parallel screening approaches require more substantial post-processing of results than combinatorial screens. Studies involving parallel screens are straightforward to design, but technical variation in how the screens are performed as well as copy number variation across cell backgrounds can confound the results (Zhang & Lu, 2009; Aguirre *et al*, 2016). Recent work has shown that copy number variation can underlie the strongest hits in CRISPR knockout screens, and multiple groups have proposed corrective algorithms to confront this problem (Pommier, 2006; Meyers *et al*, 2017; Wu *et al*). Additional heuristics aimed at increasing the quality of genetic interactions identified from parallel genetic screens have included discarding entire screens with noisy effect sizes, setting an effect size threshold for correlating genes, and capping the number of interactions per gene (Wang *et al*, 2017; Pan *et al*, 2018; Kim *et al*, 2018); however, reliance on these heuristics prevents truly unbiased genome-wide analyses. Furthermore, as the scale and diversity of published genetic screens grows, so will the need for new statistical techniques that can correct for technical variation while preserving even small levels of true signal.

In this work, we develop a flexible, unsupervised approach for removing confounding from parallel genome-wide CRISPR screens. We apply this approach to the 342 CRISPR screens of Project Achilles and compute corrected gene profiles for all reported genes. We identify more than 1 million pairs of significantly correlated, co-functional genes, substantially more than reported by other studies. Finally, we detect functionally delineated gene communities and characterize their specificity with respect to cell lineage and mutational backgrounds. Using these gene communities, we provide new insight into cancer cell network topology including scores for each gene’s potential to drive a pathway important for cancer proliferation.

## Results

### Correcting for technical confounding found in parallel genetic screen data

We first downloaded CRISPR screen gene summary data corrected for copy number confounding from the Project Achilles data depository (Meyers *et al*, 2017) and matched RNA-seq and mutation data from the Cancer Cell Line Encyclopedia (CCLE) website (Barretina *et al*, 2012). The results of the CRISPR screens form a matrix, where each row in the matrix serves as a gene essentiality profile that summarizes the knockout phenotype of the gene across the 342 cell lines (the columns of the matrix). As reported by others, the degree to which two gene essentiality profiles (rows) are correlated reflects their functional relationship. This functional relationship can reflect many gene-gene relationships, including membership in the same metabolic pathway or protein complex, and depends on the mutations present in the cell lines tested (Figure 1a, b) (Wang *et al*, 2017; Pan *et al*, 2018).

Because technical factors in the dataset could drastically skew Pearson correlation values, we explored a large set of control genes expected to have little to no phenotype in the context of cancer proliferation: olfactory receptors (as classified by the HUGO Gene Nomenclature Committee (HGNC)). Some olfactory receptors shared identical sgRNAs, and in these cases we retained only one olfactory receptor to avoid duplicate gene profiles. Within this set, all pairwise correlations were calculated under the expectation that strong effect sizes would indicate technical confounding. Under a model of uniformly null phenotypes, we would expect correlations to be tightly distributed around 0. In fact, olfactory receptors often exhibited profiles that were highly correlated across genetic backgrounds, strongly suggesting the presence of technical confounding.

To investigate the unanticipated signal of essentiality in olfactory receptors, we evaluated the covariance of the knockout growth phenotypes across cell lines using principal components analysis (PCA). We performed PCA on the matrix of growth defects from each olfactory receptor (row) and cell line background (column) (Figure 1c). The resulting principal components describe how likely each cell line is to exhibit essentiality amongst olfactory receptors. If the essentiality measurement of each olfactory receptor was being driven by gene function or chromosomal locus, these profiles – a single number per cell line – would be expected to be poorly predictive genome-wide. However, found that the top four principal components explained significantly more variance than expected by permutation testing, with over 65% of the variance explained by the first principal component (Figure S1a). We hypothesize that the reason for these significant principal components is differential cell line sensitivity to dsDNA break toxicity, something that would be well captured by a single number per cell line and consistent with prior work investigating CRISPR screen specificity (Morgens *et al*, 2017; Rosenbluh *et al*, 2017). Other cell line conditions, such as the culture media, health of the cell sample, or handling of the sample in tissue culture could also contribute technical variation captured by principal component signatures.

Having identified candidate confounding signatures, we corrected each gene’s essentiality profile by subtracting these signatures and creating an improved dataset (Figure 1c). Using these corrected gene essentiality profiles, we again computed the correlation of all pairs of genes and assessed the performance.

In many cases, as seen for peroxisome genes, learned relationships before and after correction closely resembled each other, but in others, as seen for spliceosome scaffold proteins, previously unremarkable sets of genes appeared tightly related (Figure 2a). Pairs of olfactory receptors that were persistently correlated after correction were often in very close physical proximity on chromosome arms. As observed previously (Meyers *et al*, 2017), physical proximity generally increases the rate of correlation even after CERES copy number correction (Figure S1b), suggesting either that there are substantial differences in local toxicity to dsDNA breaks or that copy number variation confounding persists at short physical distances.

**Figure 2:**
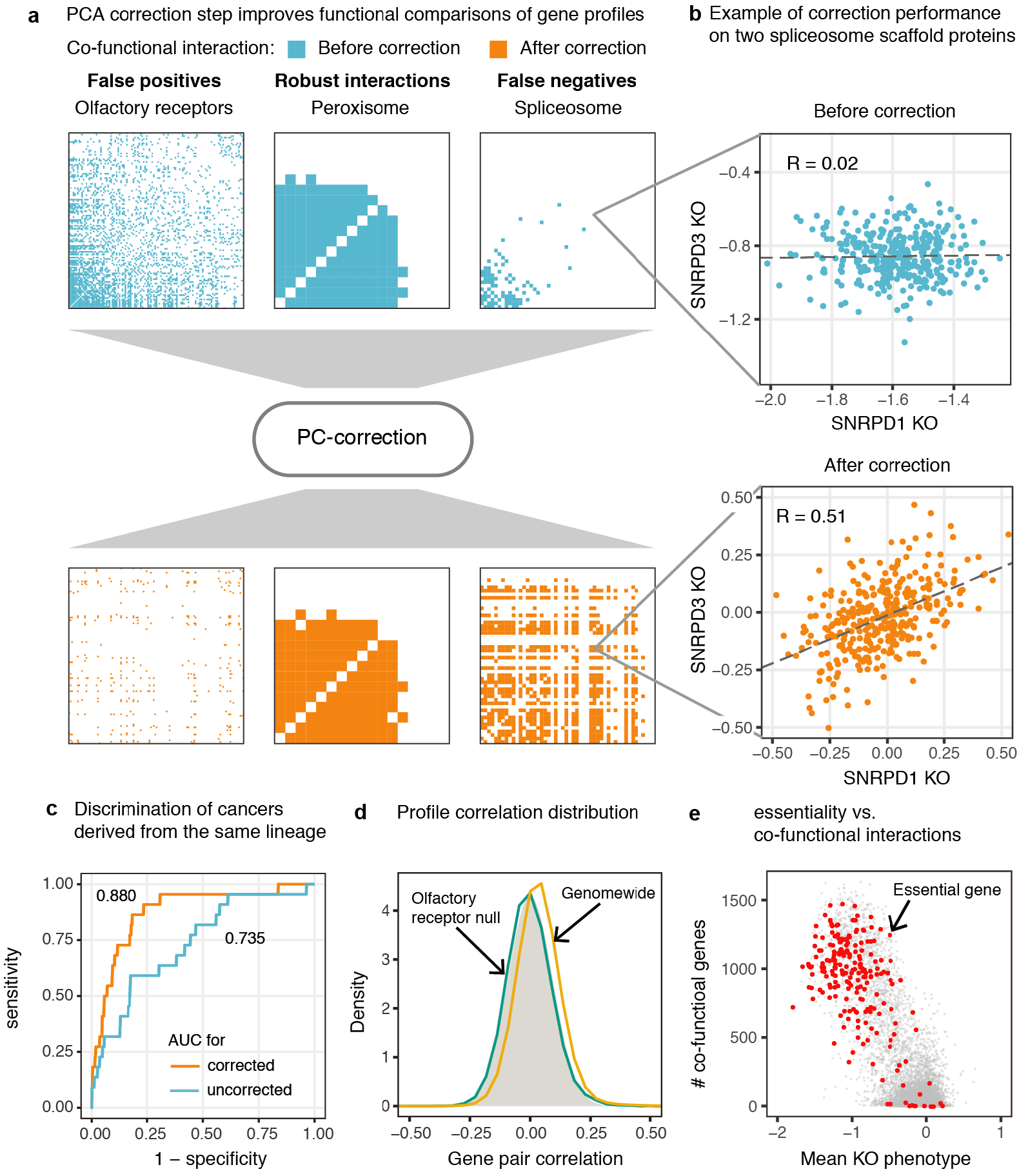
Constructing a set of co-functional genes from all pairs of human genes. **a)** Demonstration of co-functional gene calls before and after correction for three example gene sets: olfactory receptors that exhibit nonspecific interactions in the raw data, peroxisome genes that show persistent co-functionality before and after correction, and spliceosome complex members that show correlation only after correction. **b)** Correlation for two small nuclear ribonucleoproteins, *SNRPD1* and *SNRPD3*, that required for splicing, before and after correction of effect sizes from 342 cell lines. Fit lines are from linear regression. **c)** Area under the receiver operating characteristic (ROC) curve for using cell line profiles to determine if two cell lines share a developmental lineage. Cell line profiles (the vector of effect sizes across all genes for each cell line) are more correlated amongst cancers of the same type following correction. **d)** The distribution of pairwise correlations of corrected gene essentiality profiles genome-wide is greatly skewed to more positive values compared to pairs of olfactory receptors. **e)** Highly expressed essential genes regularly have on the order of hundreds of co-functional genes.

Putting aside potential confounding due to proximal genetic perturbations, we observe other clear advantages to working with corrected gene essentiality profiles. By creating cell line profiles from all gene knockout effects and comparing the correlations for every pair of cell lines before and after correction, we observed marked boosts in accuracy for predicting shared primary disease (AUC increased from 0.670 to 0.859), and for predicting shared cell lineage (AUC increased from 0.735 to 0.880) (Figure 2c, Figure S1c). At the same time the median cell line profile correlation dropped from ~0.85 to nearly 0 following correction (Figure S1d), suggesting that a single or very few axes of variation – such as relative dsDNA break toxicity – underlie large correlations between gene essentiality profiles.

### Identification of ~10^6^ co-functional interactions from corrected gene essentiality profiles

To detect gene pairs with significantly correlated gene essentiality profiles, or “co-functional” genes, we derived p-values for the observed correlations from an empirical null distribution. To build the distribution, we used the pairs of olfactory receptors described above as a background set, assuming that these receptors would not affect cancer growth. Correlations across all pairs of olfactory receptors were roughly normally distributed and centered at zero. Observed correlations across corrected gene essentiality profiles genome-wide for all other genes greatly exceeded expectation from the null model (Figure 2d). Thus, we assigned p values for every observed correlation, whether positive or negative, using a normal distribution fit to the correlations of pairs of olfactory receptors. Across all 17,670 genes tested, 1,007,278 co-functional interactions, equal to 0.65% of all possible pairs of genes, met a false discovery rate of 10%. A similar procedure on raw gene essentiality profiles yielded 17,615 interactions.

It was immediately apparent that highly expressed essential genes (Hart *et al*, 2014) generally possessed hundreds of co-functional interactions (Figure 2e). One might expect uniformly or invariantly essential genes to lack identifiable co-functional gene partners, but this was rarely the case. Furthermore, we confirmed past reports (Barretina *et al*, 2012; Hart *et al*, 2015; Wang *et al*, 2015) that genes that are either highly or invariantly expressed across tissues on average possess greater knockout effects (Figure 3a). Remarkably, this holds true not only across cell types, but even in the context of expression data from a panel of lymphoblastoid cell lines (LCLs) (Figure 3b) (Pickrell *et al*, 2010), suggesting that broad mechanisms of gene regulation and not tissue specificity are responsible for the trend. As reported previously (Wang *et al*, 2015; Pan *et al*, 2018; Wang *et al*, 2017), differences in essentiality across genes thus appear quantitative and not strictly binary across cell types. To facilitate further exploration of our co-functional gene dataset, we have developed a shiny app that visualizes all genes co-functional to the query gene with added functionality for overlaying interactions from STRING, published CRISPR screens or custom files containing scores per gene.

**Figure 3:**
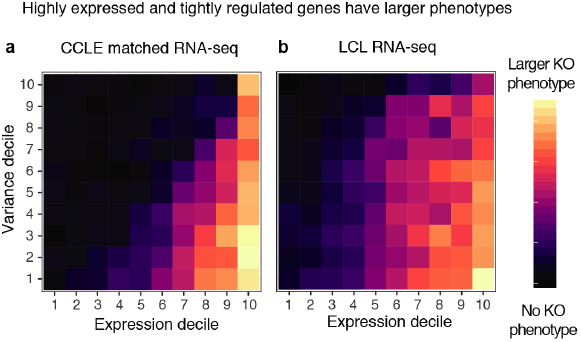
Influence of RNA expression mean and variance on gene essentiality. Genes in high mean (x-axis) and low variance (y-axis) expression bins tend to have larger knockout effect sizes, whether expression is determined from **a)** a matched RNA-seq dataset or **b)** a panel of LCL cell lines (right).

### Gene co-functionality captures variation in drug susceptibility across cancer cell lines

Past analyses (Hart *et al*, 2015; Wang *et al*, 2017) explored cell signaling in a restricted number of cancer cell lines (Figure 4a). With 342 samples, a much broader view of the diversity of signaling is possible, including differences across diverse cell types of origin. For MAPK and p53 pathways, the profiles of many genes hew closely to the *TP53* status of the cancer, producing two well-separated clusters with predictable effect sizes across cell lines (Figure 4b). Excluding a handful of outliers, *TP53* loss-of-function mutants (Barretina *et al*, 2012) do not respond to *TP53* knockout or *MDM2* knockout or treatment with Nutlin-3, an anti-*MDM2* drug. *TP53* wild type cells grow following *TP53* knockout, die following *MDM2* knockout, and exhibit slowed growth following treatment with Nutlin-3. These observations suggest that negative regulation of *TP53* is robust across cancer types, at least for its strongest co-functional genes. For example, two proposed drug targets, the deubiquitinase *USP7* (Gavory *et al*, 2018) and phosphatase *PPM1D* (Ogasawara *et al*, 2015) consistently mirror *TP53* knockout phenotypes, proving themselves robust to cell lineage and mutations in other biological pathways.

**Figure 4:**
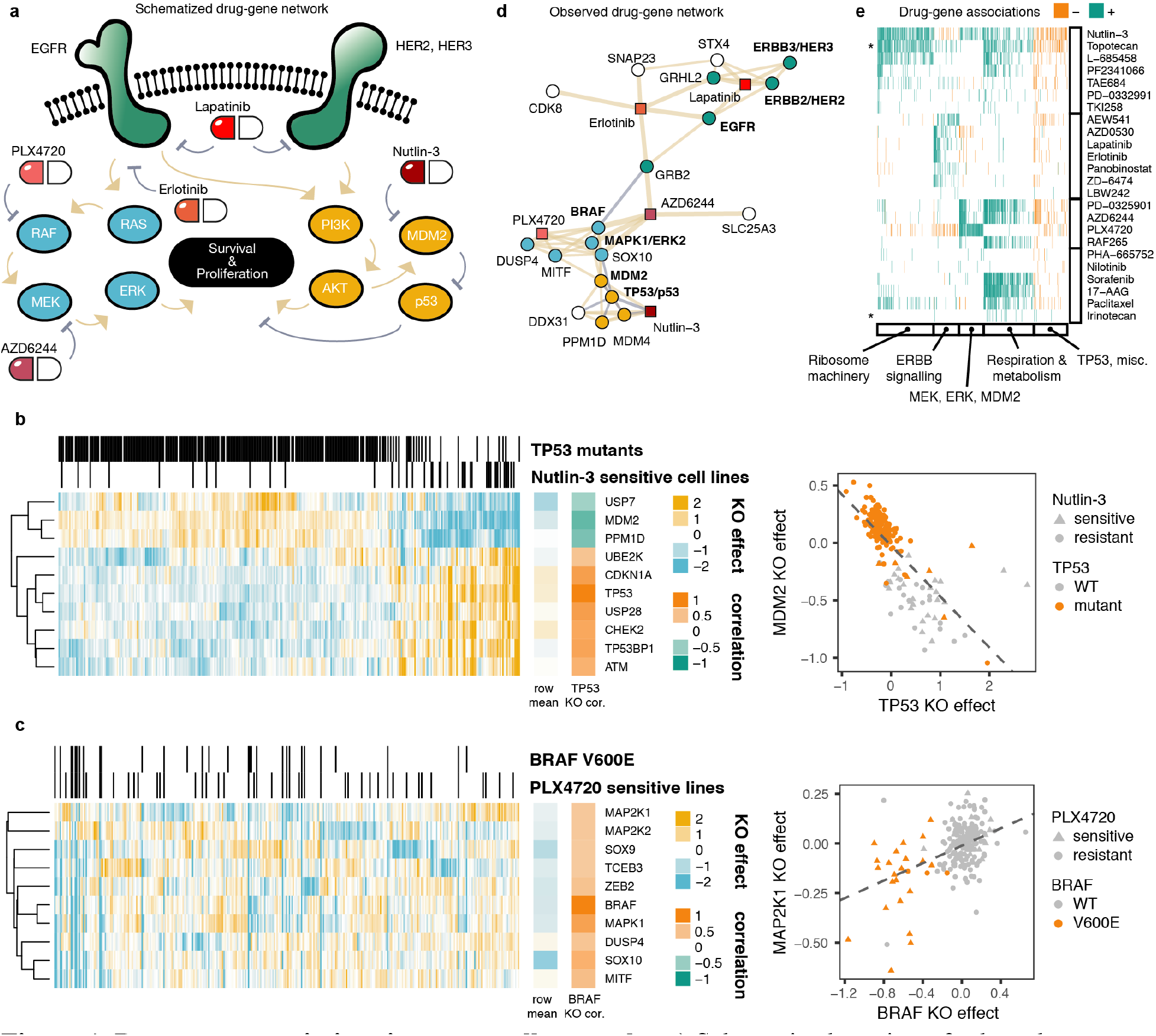
Drug-gene associations in cancer cell networks. **a)** Schematized version of selected cancer growth signaling pathways: ErbB family members (green), MAP kinase (blue) and p53-Akt (gold). **b)** Example of the most correlated genes co-functional to TP53. Each column represents one of 342 cell lines. Cell lines with TP53 loss-of-function (LOF) mutations or sensitivity to nutlin-3 (binarized by k-means) are indicated in strips above. The mean effect for each gene is shown to the right of the heatmap, as is the corrected gene essentiality profile correlation with TP53. Effect sizes are variance-normalized by row. Scatterplot of the TP53 and MDM2 rows is shown on right. **c)** Same as in **b)** but for BRAF and its top co-functional genes. **d)** Illustration of the top five drug-gene associations for each of the drugs in **a)** plus their strongest co-functional genes. Beige edges represent positive correlations, gray negative. **e)** Gene knockout effects sharing significant Pearson correlation coefficients with drug activity, clustered by k-means, for all drugs screened in the Cancer Cell Line Encyclopedia. Asterisks highlight distant clustering of topoisomerase inhibitors irinotecan and topotecan.

The essentiality of BRAF and its co-functional gene partners paints a similar picture (Figure 4c), primarily via melanoma cell lines. The knockout effects of genes co-functional to *BRAF* depend considerably on *BRAF* V600E status, with *BRAF* V600E lines especially sensitive to *BRAF*, *MAP2K1* and *MAPK1* knockout. Our findings are similar to those independently reported by Kim, et al when this manuscript was in preparation. Thus, there is strong evidence that genes co-functional to central cancer growth genes mediate their essentiality through their involvement in those genes’ pathways.

### Incorporating drug-gene associations into gene networks

By correlating the maximal activities of anticancer compounds in each cell line to the gene profiles derived from knockout effects, it is possible to test for drug-gene associations and combine drugs and genes into a single network (Figure 4e, Figure S2). The ErbB family of proteins includes four well-characterized receptor tyrosine kinases (RTKs) that act upstream of PI3-K and MAPK pathways and complex with each other to mediate growth signaling (Medina & Goodin, 2008). ErbB family members are highly druggable and have been targeted by erlotinib, an anti-EGFR drug, and lapatinib, a dual anti-HER2, anti-EGFR drug. We confirmed that lapatinib-sensitive cell lines were typically sensitive to knockout of ErbB family members (*EGFR*, *ERBB2*/HER2 and *ERBB3*/HER3), whereas erlotinib sensitive lines were associated with sensitivity to *EGFR* and not other ErbB members. The important role of *GRB2* in mediating *EGFR* signaling to the MAPK pathway is shown as drug-gene associations linking *GRB2* to both erlotinib and AZD6244, a MEK inhibitor. Somewhat counterintuitively, *GRB2* knockout sensitivity is negatively correlated with *BRAF* knockout sensitivity because the most *BRAF*-dependent lines (*BRAF* V600E melanoma) are not sensitive to *EGFR* knockout, producing a mutually exclusive relationship.

After pooling all drug sensitivity data, drug-gene associations clustered into approximately five classes (Figure 4e). Foremost among these was a large number of gene knockouts enriched for terms relating to ribosome biogenesis and DNA break repair that were correlated with sensitivity to nutlin-3. Because cell stress rapidly induces p53 signaling (Zhang & Lu, 2009), we might expect gene knockouts that interfere with ribosome activity or DNA repair to probe the integrity of the p53 pathway: if a cell stress-inducing knockout swiftly kills a cell line, it is likely that p53 is intact in that cell line. Interestingly, topotecan participates in many more drug-gene associations with these genes than the mechanistically related drug irinotecan (Pommier, 2006). The second class of genes exhibits interactions for the same drugs as the first but features mostly negative drug-gene associations and is unremarkable except for containing *TP53*.

The third class of drug-gene associations most notably features numerous ErbB signaling genes. In general, knockout of these genes is prognostic for sensitivity to receptor tyrosine kinase inhibitors like erlotinib, lapatinib, ZD6474 and AEW541. A fourth class is mostly driven by MEK and ERK signaling in melanoma cell lines dependent on *BRAF* V600E. These lines are uniquely susceptible to PLX4720. In fact, the mutual exclusivity of PLX4720 sensitivity and ErbB knockout sensitivity produces several negative drug-gene associations between the cluster of ErbB signaling genes and PLX4720.

A final class uncovered a link between perturbation of respiration and metabolism genes and broad sensitization to many different drugs. To explain this, we hypothesize that cell lines that are sensitive to metabolic perturbation are, on average, more likely to be susceptible to pharmacological perturbation.

### Enrichment of co-functional genes in curated gene set databases

To evaluate the extent to which co-functional genes reflect distinct kinds of functional relationships, we calculated the enrichment of co-functional gene relationships across gene sets maintained by the Molecular Signatures Database (Subramanian *et al*, 2005). Globally, curated gene sets contained more co-functional pairs of genes than expected by chance: two-fold enrichment for genes annotated with the same biological process or molecular function, more than threefold for cellular component, and more than fivefold for KEGG pathways (Figure 5a) (Ashburner *et al*, 2000; The Gene Ontology Consortium, 2017). With respect to reconstituting curated gene sets, our co-functional gene dataset exhibits lower enrichment than state-of-the-art protein mass spectrometry but contains 70 times more potential interactions (Rolland *et al*, 2014). Even with low enrichment, approximately one-third of biological process and 60% of cellular component terms were enriched above random matched networks (Figure 5b, see methods). We also performed gene set enrichment on genes with no co-functional gene edges and found that underrepresented pathways were mostly limited to genes involved in lineage-specific (e.g., muscle) development and differentiation (Figure S3a). Despite these blindspots, we conclude that many diverse biological pathways, not simply pathways typically associated with essentiality, contribute to gene co-functionality.

**Figure 5:**
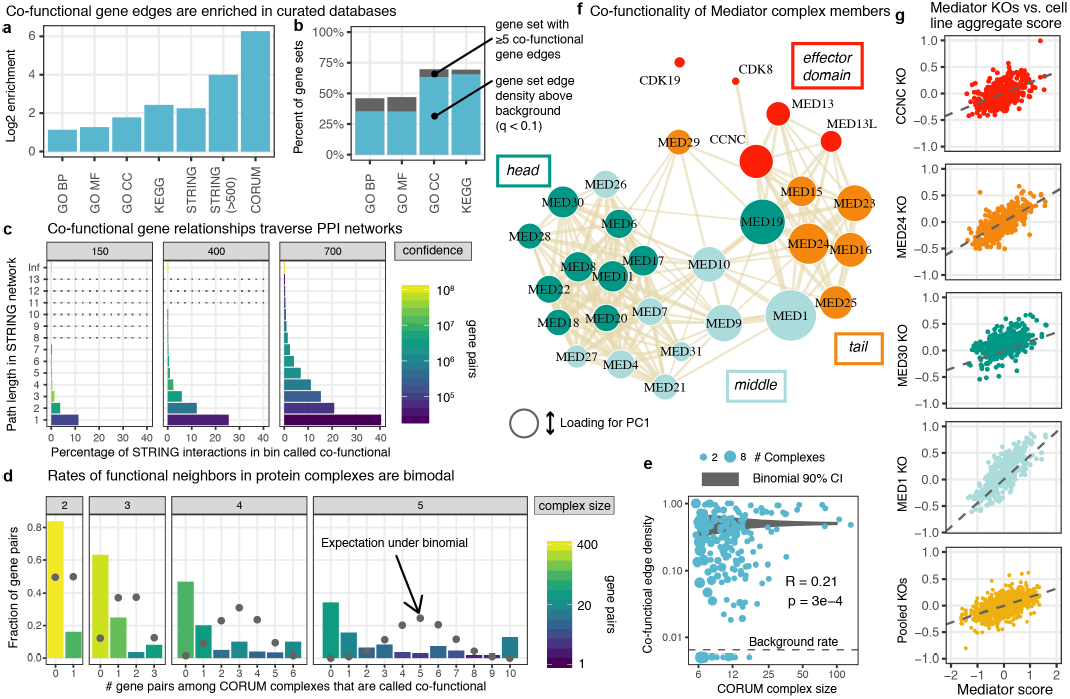
Comparison of gene co-functionality to diverse gene annotation databases. **a)** Pairs of genes annotated with the same Gene Ontology term (BP = biological process, MF = molecular function, CC = cellular component) or Kyoto Encyclopedia of Genes and Genomes (KEGG) pathway are enriched for co-functional interactions. Pairs of genes with annotated interactions from STRING, especially high confidence (>500) interactions, are also enriched. Pairs of genes belonging to the same protein complex as curated in the Comprehensive Resource of Mammalian Protein Complex (CORUM) core complex database exhibit the greatest enrichment. **b)** Percent of gene sets containing at least 5 pairs of co-functional genes and, amongst those, the fraction that are enriched for co-functional interactions above degree-matched random graphs. **c)** Rates of co-functional gene calls for gene pairs binned by their path length in the STRING experimentally derived PPI network. “Inf” (infinite) refers to genes that are in separate components. **d)** Co-functional call rates amongst CORUM protein complex edges split by complex size. The binomial expectation for the number of edges called as co-functional is shown in gray. Co-functionality rates per complex are bimodal and greater for large protein complexes. **e)** Extension of **d)** to larger protein complexes, with the number of edges expressed as the fraction of all possible edges in the complex, equal to 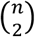. The area of each point reflects the number of multiple protein complexes with the same complex size and edge density. Complexes with fewer co-functional genes than expected according to the average rate are shown below the dashed line. The 90% binomial confidence interval for random co-functionality calls given the average rate is shown in gray. Large complexes again exhibit more co-functional interactions. **f)** Reconstruction of the Mediator complex from gene co-functionality, shown in a force-directed graph. Subunits in the same domain (head, middle, tail, or CDK effector) of the complex are more likely to be co-functional than subunits in different domains. The area of each gene node corresponds to its loading on the first principal component of the corrected gene essentiality profile matrix. **g)** Mediator subunit knockout effect sizes in each cell line can be summarized by a “Mediator score,” the first principal component score for each cell line, plotted against the knockout phenotype of specific genes (first four facets) or a pool of all genes in the complex (bottom row).

### Co-functional gene edges overrepresented amongst protein-protein interactions

Relative to curated gene sets, reported protein-protein interactions exhibited greater enrichment of co-functional genes. This was true for large-scale repositories of experimental data, as curated by the STRING consortium (Szklarczyk *et al*, 2015) (16-fold amongst interactions above 500 confidence), and for well-studied protein complexes, as curated in the CORUM core complex resource (Ruepp *et al*, 2010) (77-fold for proteins in the same complex). In fact, we identified the majority (50.2%) of gene pairs in CORUM core complexes (N = 48,408) as co-functional genes (binomial test p < 1e-300).

We next explored how STRING interactions related to our set of co-functional genes in more detail. 14% of co-functional interactions were present in the STRING database of protein-protein interactions, compared to 3% of all pairs of genes. The probability of a STRING interaction being called varied depending on the type of annotation and in accordance with its confidence (Figure S3b). 7% of low confidence (score 1-150) experimentally derived STRING interactions were amongst our co-functional interactions, compared to a full 82% of high confidence (score 900-1000) STRING interactions. Likewise, only 3% of low confidence co-expression transferred STRING interactions were amongst our co-functional interactions, compared to 64% of high confidence STRING interactions. Nonetheless, all categories were enriched for co-functional genes above the global average of 0.65%.

Interestingly, we find only a small fraction (~3%) of co-functional genes to be negatively correlated. Amongst pairs of genes in STRING with detailed functional annotations, 21% of the highest confidence (>900) activating interactions were called, in contrast to 7.3% of highest confidence inhibiting interactions (Figure S3c). High confidence (score > 700) activation, catalysis, reaction and binding interactions were overwhelmingly positively correlated (99.0%, 99.7%, 99.9% and 99.9% of correlations above zero, respectively, Figure S3d). While the statistically significant correlations for inhibiting interactions might be expected to be broadly negative, this was not the case: only 2.7% of correlations were negative. We found that 64% of STRING interactions annotated as inhibiting were also annotated as activating or catalyzing and could predominantly operate in a cooperative rather than inhibitory manner, but even among pairs of genes exclusively annotated as sharing an inhibitory relationship, only 5.8% were negatively correlated. This dearth of anti-correlated co-functional genes is consistent with the observation by Wang, et al. that anti-correlated co-essential genes are rarely detected (Wang *et al*, 2017). Pooled CRISPR knockout screens, even with diverse panels of cell lines, may have low sensitivity for detecting inhibitory relationships across pairs of genes, or negative regulation more broadly.

Because our reported number of co-functional interactions exceeds the estimated number of human gene pairs that physically interact (Venkatesan *et al*, 2009; Stumpf *et al*, 2008) by as much as tenfold, we explored the potential underpinnings of co-functionality in greater detail. While we infer a substantial fraction of experimentally validated genetic interactions in STRING as co-functional, most of our co-functional gene pairs fail to appear STRING’s database of experimentally supported interactions. In total, only 13% of co-functional genes are annotated by any co-expression/experimental study, suggesting that physical interactions among proteins and gene co-regulation explain a minority of co-functional interactions. To follow up on this result, we calculated the rate of gene co-functionality as a function of the path length dictated by the STRING database to evaluate the extent to which co-functional genes could be linked indirectly by cascades through the human interactome. Pairs of genes separated by two to four experimentally derived STRING edges had consistently higher rates compared to pairs of genes in separate components of the network. Amongst STRING interactions of confidence > 700, genes with one intermediate gene, or path length 2, were enriched 51-fold (Figure 5c). In addition to gene regulatory and metabolic relationships that do not require physical interaction, signaling cascades that travel via multiple binary interactions can underlie additional co-functional interactions.

### Characterizing co-functionality amongst members of the same protein complexes

Across CORUM core complexes, we note that the number of co-functional interactions per complex deviates far from expected assuming independent draws from a binomial distribution. In fact, oftentimes all or none of a protein complex’s members were labeled co-functional (Figure 5d, e). Complex size also played a role, as 67% of core complexes with 5 or fewer members contained no co-functional genes, whereas 91% of complexes with more than five members exceeded the average rate of co-functional interaction calls. Protein complex essentiality appears to explain some of this bimodality: co-functionality is rarely detected amongst CORUM core complex members that only weakly affect growth, whereas CORUM core complexes that strongly affect growth are almost complete graphs of co-functionality (Figure S3e), as seen for mitochondrial and cytosolic ribosomes, the proteasome, U2 snRNP, and RNA polymerase II complexes.

Although the rate at which we identified the members of a protein complex co-functional varied according to the knockout growth phenotype, we identified other factors that guided which of a complex’s members were co-essential and which were not. The Mediator complex, a transcriptional coactivator that has been previously examined using a similar parallel screening analytical approach (Pan *et al*, 2018), serves as an example. Although knockout of every Mediator complex member is associated with impaired growth, we called fewer than half of the pairs of genes comprising the Mediator complex co-functional (Figure 5f). The network of co-functional gene edges describing the members of the Mediator complex in fact mirrors the 3-dimensional structure and function of the complex; gene knockouts cluster by their topology, divided into head, middle, tail and effector module (Yin & Wang, 2014). We also summarized cell lines by running PCA on the matrix of cell lines by Mediator complex members and assigning the first PC score of each cell line as its “Mediator score”. This score integrates the 30 Mediator member knockout effect sizes into a single number reflecting the growth defect associated with Mediator loss of function. The Mediator score can predict the effect of knocking out arbitrary Mediator complex members (Figure 5g) but does so independently of protein domain-specific effects.

### Gene communities in the cancer cell network offer insight into cancer proliferative processes

To examine the distribution of co-functionality genome-wide and nominate candidate core genes for cancer growth, we performed community detection to partition all genes into separate communities (Fortunato, 2010). Every gene was assigned to one of 2,857 separate communities, including 2,139 singleton genes with no co-functional genes and 487 small communities with fewer than 8 genes. Because singleton and very small gene communities are information-poor and harbor few edges, we focused on the 231 communities with at least 8 gene members (Figure 6a).

**Figure 6:**
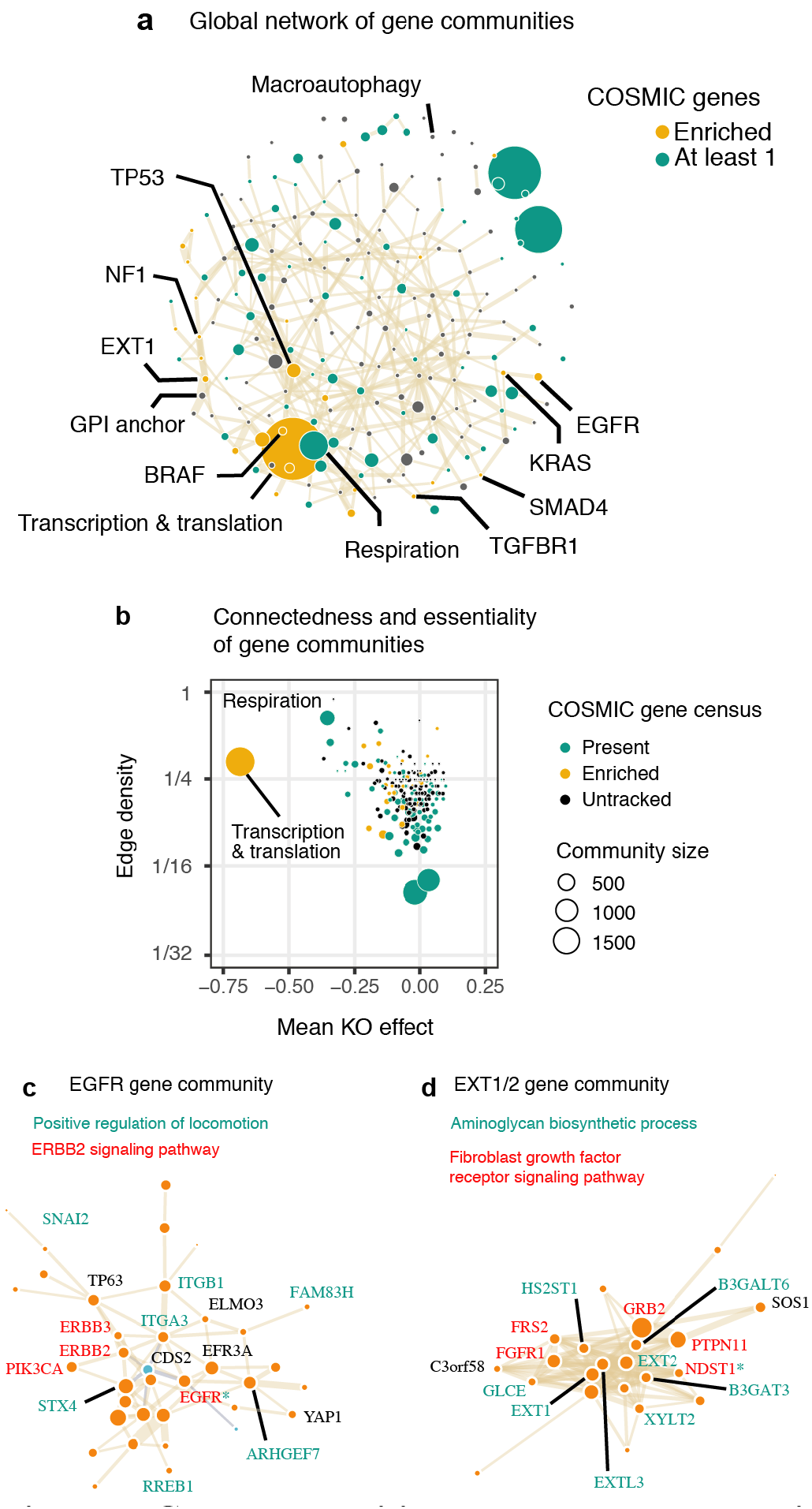
Gene communities and network topology in cancer cell networks. **a)** Consolidation of genes into >200 communities using de novo community detection. COSMIC census genes frequently cluster in the same gene communities beyond expected by chance (in gold) but are otherwise widely dispersed throughout the network (in green). Nodes representing communities are sized by number of constituent genes. The largest community contains over 1000 core essential genes (labeled ‘Transcription & translation’). Select gene communities are labeled by enriched annotations and/or prominent cancer-associated genes. **b)** Distribution of gene communities by mean knockout effect size of community members and co-functional edge density. ‘Respiration’ and ‘Transcription & translation’, from **a)** are both broadly essential and densely connected. **c-d)** Depictions of the *EGFR* and *EXT1* gene communities. Size of the gene nodes reflects loading on the first principal component amongst community members. Gene symbols that are members of selected enriched gene ontology terms are labeled.

We first enumerated the 719 COSMIC census genes in each community (Forbes *et al*, 2017) and found that the majority of gene communities (118 / 231) contained at least one census gene, illustrating the extreme diversity of pathways that cancers can manipulate to maximize cellular proliferation. However, certain gene communities possessed many more census genes than expected, including communities containing *TP53*, *EGFR*, *KRAS*, and *BRAF*. More than half of the clusters (117 / 231) were significantly enriched for at least one Gene Ontology term or KEGG pathway at a false discovery rate of 10%.

The largest community we identified is densely connected (33% of all pairs connected) and encompasses numerous core essential processes identified by KEGG, including the spliceosome, the ribosome, the proteasome, cell cycle, RNA polymerase, and mismatch repair (Figure 6b). A smaller, even denser cluster (67% of all pairs connected) contains genes required for mitochondrial function and aerobic respiration. Genes lacking any pathway annotations for the most part were not enriched in any gene communities but were significantly dis-enriched in these core growth communities (Figure S4).

Gene communities varied substantially in the breadth of their encapsulated functions. Some communities derived from specific cancer pathways, such as the community containing *EGFR*, which simultaneously captured signaling from related ErbB proteins and the role of cell adhesion and locomotion in *EGFR*-dependent transformation (Figure 6c) (Lindsey & Langhans, 2015). Other clusters reflected very specific cell functions or compartments, as in the case of the peroxisome.

With respect to the peroxisome community, 29 Gene Ontology terms were statistically enriched, ranging in size from 13 to 406 genes. Enriched gene sets primarily derived from peroxisomal transport and fatty acid metabolism. Although these are not conceptually similar pathways, they both relate to the core function of peroxisomes, which are variably essential across the cancer cell lines. In the context of cancer proliferation, abstracting fatty acid metabolism away from peroxisomal transport genes, as done in the Gene Ontology database, may ignore the reality that these processes are functionally inseparable.

Another community (Figure 6d) contains the genes underlying hereditary multiple osteochondromas, a rare Mendelian disorder. These genes, *EXT1* and *EXT2*, are known to act in the polymerization of the glycosaminoglycan heparan sulfate, a known cofactor for FGF signaling (Ornitz & Itoh, 2015). How *EXT1* loss leads to cancer is poorly understood (Bovée, 2008); nonetheless, eight other known aminoglycan synthesis genes participate in the same gene community. Also present are fibroblast growth signaling genes *FGFR1*, *FRS2*, *GRB2*, *NDST1* and *PTPN11*. One possibility is that every member of the aminoglycan synthesis pathway influences cancer proliferation by ultimately making heparan sulfate available to upregulate FGF pathways. If true, the number of genes that could potentially modify cancer proliferation via FGF signaling is much larger than currently appreciated, expanding from *EXT1* and *EXT2* to all aminoglycan synthesis genes.

### Gene network centrality adds another layer to gene function

We next examined the network topology within and across gene communities. Because core growth genes would dominate strength or degree calculations for most genes, we quantified the centrality of every gene using the closeness of each gene within its prescribed community. The closeness of a gene is the reciprocal of the sum of the shortest distance via co-functional gene edges from that gene to every other gene in the network. By calculating this metric within communities, we attain a local centrality measure that gives insight into a diverse range of gene functions.

As a measure of centrality in the cell network, closeness added information not captured by other gene properties such as essentiality or expression. Overall, COSMIC census genes did not differ in closeness compared to other genes (Wilcoxon rank-sum p > 0.05). Neither did they differ substantially in their average knockout growth phenotype. However, genes annotated as germline cancer genes scored much higher in closeness than somatic cancer genes (Figure 7a, Wilcoxon rank-sum p = 7e-9). In fact, gene closeness surpassed gene expression level or knockout growth phenotype in accuracy for ascertaining somatic from germline cancer genes (Figure 7b). Closeness was slightly greater for genes suspected to be subject to strong purifying selection via population genetic data (high pLI genes) (Lek *et al*, 2016), but high RNA expression level was more predictive of high pLI status than closeness (Figure S5a). We also observed that genes linked to unfavorable prognoses in cancer patients exhibited greater closeness in gene communities, but that genes linked to favorable prognoses did not (Figure 7c) (Uhlen *et al*, 2017). In all, we linked high closeness genes to oncogenes and germline cancer processes, complementing broad patterns of high expression and essentiality for cancer drivers.

**Figure 7:**
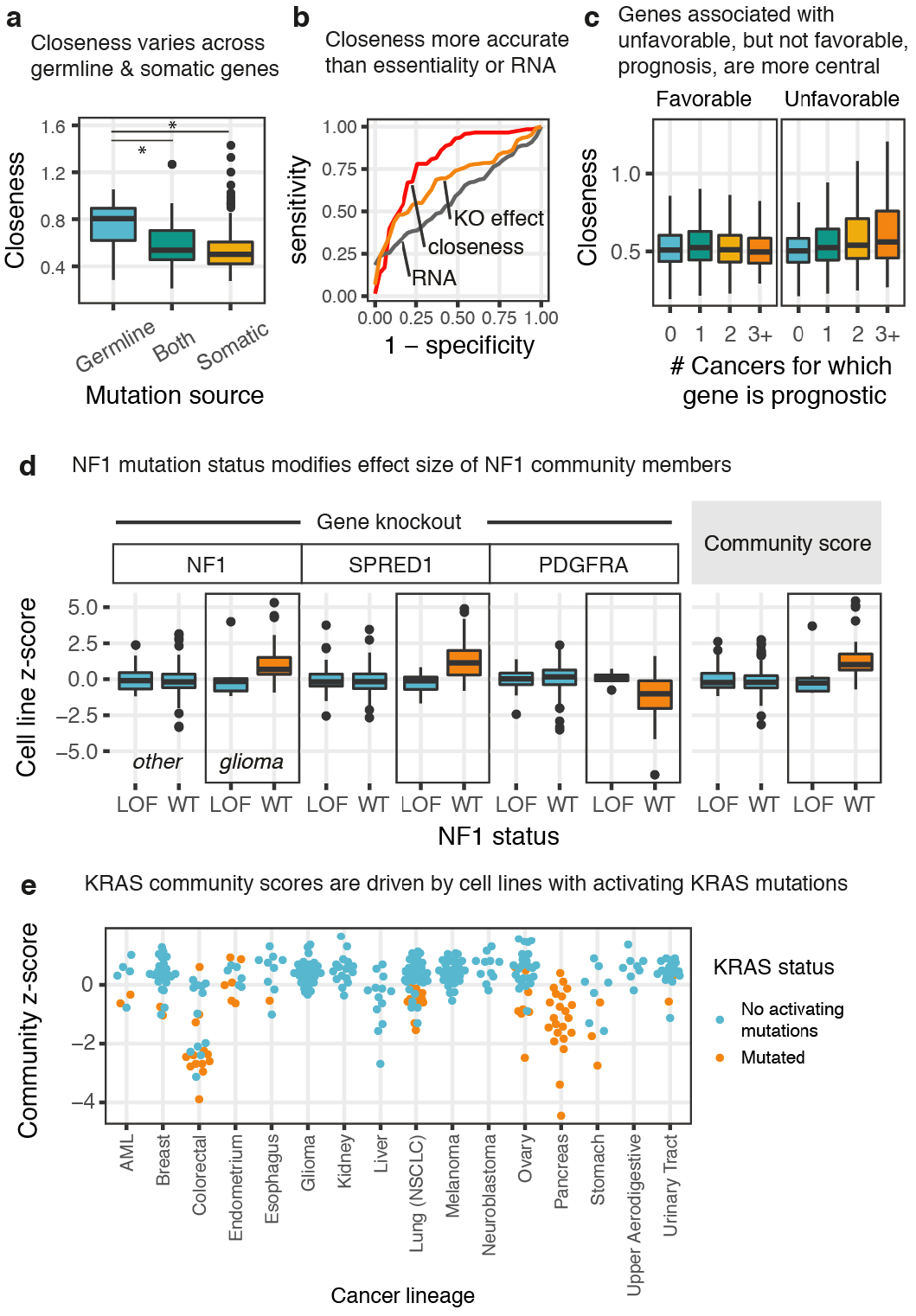
Topology and cell type specificity of cancer networks. **a)** Boxplot of closeness for genes annotated as germline cancer genes, somatic cancer genes, or both. **b)** ROC curve showing how centrality (measured by closeness in the gene community) separates germline from somatic cancer genes in the COSMIC gene census more accurately than either growth knockout phenotype or expression. **c)** Genes associated with unfavorable prognoses as found in the Human Protein Atlas Pathology database have greater closeness within gene communities, left. The same is not true for genes associated with favorable prognoses, right. **d)** NF1 gene community knockout phenotypes and the NF1 aggregate community score calculated from the first principal component of member genes shows that perturbation of the community by CRISPR knockout almost always occurs in NF1-WT glioblastoma cell lines but only rarely in other cancers. P < 0.05 for all within-glioma comparisons. **e)** KRAS community scores demonstrate that dependence on constitutively active (mutant) KRAS drives community organization, independent of cell lineage, but is uniformly present across pancreatic cancers.

To summarize cancers by gene community activity, we again performed principal components analysis – this time separately for each gene community (see methods) – and recorded the first PC score for each cell line. The first PC score reduces the dimensionality of the gene profiles from the size of the gene community down to a single number, which we term the community score. A potential benefit of scoring communities as opposed to individual genes is that the shared function of a gene community can be ascertained while avoiding the measurement noise or alternate signals associated with individual gene knockouts.

We first explored how community score varied by cell lineage and confirmed that cancer cell lineages dependent on particular growth pathways were associated with extreme scores for the corresponding gene communities. For example, it was often possible to classify the lineage of glioblastoma, neuroblastoma, small cell lung cancer, melanoma, and pancreatic- and kidney-derived cancers by virtue of their especially extreme community scores (Figure S5b), a finding also reported by Kim, et al. Interestingly, we found that communities associated with cell lineage identity (see methods) were five times more likely to be enriched for COSMIC census genes than gene communities with no associations (Fisher exact test p = 4e-6), consistent with the interpretation that cancers from different cell lineages target different central growth pathways to maximize growth.

### Diversity of cell line panel expands the scope of detectable cell signaling paradigms

In some cases, cell lines of a specific lineage and mutational background are required to uncover cancer-relevant growth pathways. The gene community containing the tumor suppressors *NF1* and *SPRED1* illustrates this phenomenon (Figure S5c). In the germline, one *NF1* loss of function allele causes the Mendelian disorder Neurofibromatosis type 1 (Gutmann *et al*, 2017). Similarly, loss of function of *SPRED1* causes Legius syndrome, which can be confused clinically with Neurofibromatosis type 1 (both genes act upstream of RAS signaling). In the panel of cell lines from Project Achilles, glioma cell lines consistently exhibit the largest effect sizes for community members *NF1*, *SPRED1* and *PDGFRA*. Yet, even amongst glioma cell lines, only those that lack a pre-existing *NF1* loss-of-function event exhibit large differences in sensitivity to gene knockout of community members (Figure 6d). The differences across these genes are well summarized by *NF1* community scores, where glioma cell lines with intact *NF1* consistently score above glioma cell lines with *NF1* loss of function (Wilcoxon rank-sum p = 0.005). In the case of non-glioma cell lines, NF1 community scores rarely deviate from the mean and do not vary by *NF1* mutation status (Wilcoxon rank-sum p > 0.05).

While extreme community scores often occur in distinct cell lineages, sensitivity to perturbation of gene communities can often be accessed across multiple lineages. For example, the *KRAS* gene community is strongly associated with pancreatic and colorectal cancers, but also exhibits apparent activity in cell lines derived from other lineages. In general, it is apparent that extreme *KRAS* community scores indicate the presence of mutant, constitutively active *KRAS* (Figure 7e), with pancreatic and colorectal cancer very likely to acquire such mutations. In the Project Achilles panel, all pancreatic cancers harbor *KRAS*-activating mutations, while ovarian cancers rarely do. This difference in mutational status explains the difference in *KRAS* community scores between pancreatic and ovarian cancers more accurately than cell lineage and stands in contrast to the *NF1* gene community example where both a specific lineage and mutational status were required.

## Discussion

This work demonstrates the power of unsupervised statistical techniques for correcting gene profiles constructed from parallel screening datasets. We show that technical confounding is pervasive across CRISPR knockout screens, but that highly active sgRNA libraries and extensive data preprocessing steps can expand the quantity and quality of interactions called from correlated gene profiles, whether or not interactions are mediated directly by protein-protein interactions (Figure 8). When screening for hit genes that modify a phenotype of interest, it is already considered best practice for sgRNA libraries to contain “safe-targeting” sgRNAs that target non-genic regions to correct for the toxic effects of DNA cleavage (Morgens *et al*, 2017). We provide evidence that sgRNA off-target effects can cause both false positive and false negative ascertaining genetic interactions, and, by exploiting a set of control genes with a very low prior of having an effect on the screen phenotype, we can correct these confounding signatures. In the future, more comprehensive sets of control sgRNAs may further improve modeling of confounding across genomic regions of variable copy number, sequence content and chromatin accessibility.

**Figure 8:**
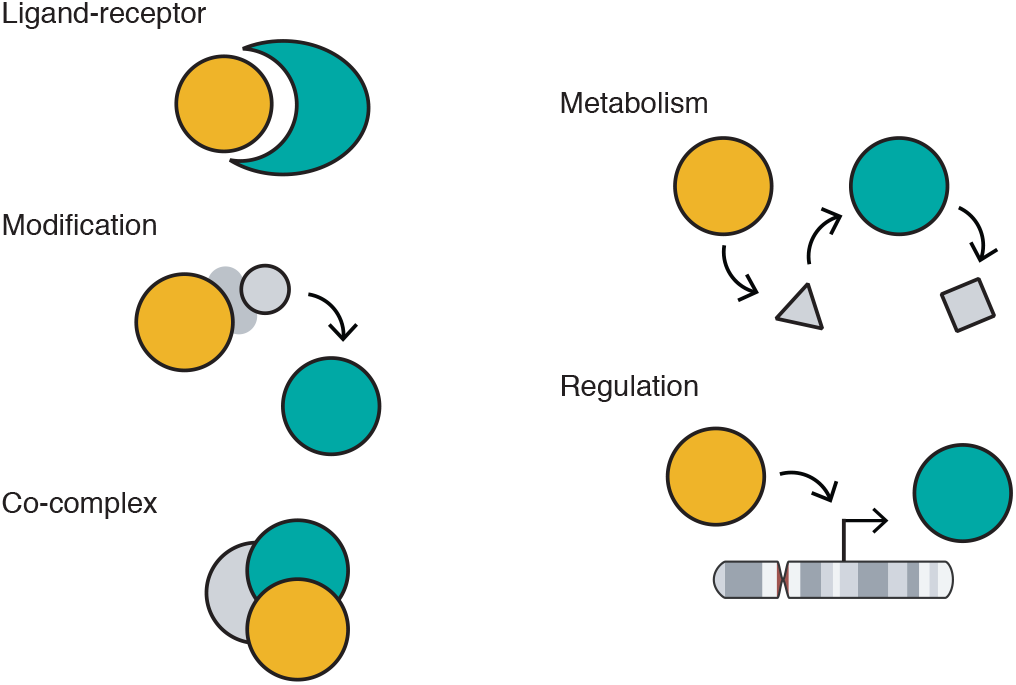
Co-functional gene relationships. Co-functionality learned from correlations between corrected gene essentiality profiles broadly extends to gene-gene relationships from physical interactions like ligand-receptor interactions, co-complex formation, or post-translational modification to conceptual relationships like gene regulation and shared metabolic pathways.

Although some biological pathways can be readily detected using a relatively small number of genetic screens for cancer growth, a comprehensive mapping of cell networks will require much more diverse panels of screens. In multiple cases, we found cancer pathways that were active only in specific cell lineages or mutational backgrounds. Amongst screens currently available, inhibitory relationships among genes are only rarely detected. Finally, gene knockouts that lack a growth phenotype in the available cell lines cannot be incorporated into cell networks at all. These findings argue for collecting screen data for samples of cells with diverse mutational and cell lineage backgrounds and across diverse screening conditions, including activating (CRISPRa) screens that might improve detection of antagonistic gene pairs. Parallel screens in the same cell lines for cancer phenotypes other than growth, such as invasiveness or cell size, would complement existing data on proliferation, but screening other phenotypes, such as phagocytosis or sensitivity to oxidative stress, offers a way to improve the richness of sparse gene profiles. As more screening data become available that differ by screening phenotype, lab, manner of perturbation (CRISPRi/a, base editing, inducible systems), and sgRNA library, quality control for parallel screening techniques will become increasingly critical.

Maps of cancer cell networks drawn using co-functional interactions are assuredly detailed, but parallel screening approaches come with distinct limitations. First, the co-functional interactions we report obscure the distinction between direct and indirect interactions as well as the directionality of information flow in signaling processes. De novo methods to call gene communities might cluster genes into a smaller set of modules, but the manner by which genes in the same module work together remains difficult to infer. One open question specific to the field of parallel screening is whether interactions identified by double knockout screens, particularly synthetic interactions, can be predicted from co-functional interactions identified in data collected from other cell lines. Also unknown is the extent to which gene communities learned from one phenotype are the same communities underlying other phenotypes. Yet, information theoretic models and orthogonal genome-wide profiling data (RNA-seq, ATAC-seq) promise to expand what can be learned from genome-wide perturbation data. In spite of their limitations, genome-wide perturbations remain a powerful tool for comprehensively unpacking the logic of cell networks.

## Materials and Methods

### Calling co-functional interactions from CRISPR screening data

Project Achilles gene level effect sizes, the list of 217 highly expressed genes from Hart, et al., and sgRNA sequences were downloaded from Meyers et al.’s data depository (https://figshare.com/articles/_/5520160) for further processing.

To correct cell line-specific Cas9 toxicity, nonspecific cell line signatures were generated from olfactory receptor gene essentiality profiles. The HGNC olfactory receptor gene list was downloaded from the HGNC website (https://www.genenames.org/cgi-bin/genefamilies/set/141). The full list of olfactory receptor gene symbols was intersected with gene symbols present in the gene level statistics. Some sgRNA sequences were not unique amongst olfactory receptors; in these cases, one olfactory receptor was selected at random and any olfactory receptors with duplicate sgRNA were discarded. The resulting matrix of 250 olfactory receptors by 342 cell lines was transposed and subjected to PCA using the *prcomp* function in R. The same procedure was applied to 100 permutations of the same matrix by shuffling effect sizes within columns (cell lines). For the first four PCs, the proportion of variance explained per PC for the true matrix exceeded that of all permutations and were deemed signatures of nonspecific toxicity.

Gene essentiality profiles were projected onto the principal components identified as nonspecific and converted back to the original dimension via matrix multiplication, and the difference of matrices was taken as a set of corrected gene essentiality profiles.

In contrast to the original, frequently positively correlated gene essentiality profiles, correlations among corrected gene essentiality profiles for olfactory receptors resembled a normal distribution centered at zero. A null distribution was created with mean 0 and standard deviation equal to the square root of the mean squared effect size (Figure 2d). P-values were assigned to all human gene pairs using this normal distribution via the *pnorm* function in R, and hits at a 10% FDR were identified as co-functional interactions.

Using the same correlation cutoff for gene essentiality profiles, co-essential gene hits were compared to co-functional genes within three sets of genes: Gene Ontology ‘spliceosomal complex assembly’ genes (a category with surprisingly few correlated gene essentiality profiles), Gene Ontology ‘peroxisomal transport’ genes (a well circumscribed set of co-essential genes) and olfactory receptor genes with co-functional degree of 15 or greater. In general, the number of co-functional interactions, or degree, varied considerably across all genes. Highly expressed essential genes were confirmed as being greatly enriched among genes with very high degree (Figure 2e).

### Predicting shared lineage and primary disease from cell line signatures

Cell line profiles were taken from the transpose of the corrected gene essentiality profile matrix and pairs of cell lines were labeled as deriving or not deriving from the same cell lineage. This created 342 choose 2 observations of 0 (different lineage) and 1 (same lineage). The same was done for primary disease. ROC plots and AUC values on using correlation between cell line profiles to detect lineage or primary disease identity were calculated using the ROCR R package. The higher the accuracy, the better gene profiles are able to expose differences across cell types in biological pathway dependence.

### TP53 and BRAF co-functionality heatmaps

Germline-filtered mutation data, including for TP53 and BRAF, and drug sensitivity data was downloaded from the CCLE portal (https://portals.broadinstitute.org/ccle/data). For TP53, all cell lines with a protein-coding or splice mutation were labeled loss of function mutants. For BRAF, cell lines with a V600E protein change were labeled V600E lines. Binarized drug sensitivity to either Nutlin-3 or PLX4720 followed from k-means clustering on the Amax values with k set to 2 to separate two groups of resistant and sensitive.

The top 9 genes co-functional to either TP53 or BRAF were visualized in the heatmap using the R package ComplexHeatmap. Corrected gene essentiality profiles (rows) were divided by their corresponding standard deviation to normalize the heat scale. The distance metric for genes (rows) was magnitude of Pearson correlation. For cell lines (columns), signed Pearson correlation was used. Accompanying scatterplots show the unnormalized gene knockout effect sizes with a linear fit trend line.

### Drug-gene network and associations

The reported Amax values were used as measures of drug efficacy for all cell lines. Drug phenotypes were then correlated with gene knockout effect sizes for all drugs across all genes. For network visualization (Figure 4d), the four genes with the largest magnitude correlation in either direction were selected. For all drug-gene associations (Figure 4e), hits were called for correlation p-values meeting a 10% FDR cutoff. Amongst these, a positive correlation between a gene and a drug denotes that cell line growth impairment after gene knockout is associated with greater growth impairment upon drug treatment. This is expected when the same oncogenic pathway is targeted by the drug as by then gene knockout. A negative correlation denotes that cell line growth following gene knockout is associated with greater growth impairment upon drug treatment. This is expected for tumor suppressor-drug pairs wherein the pathway the drug inhibits is negatively regulated by the tumor suppressor, as with Nutlin-3 and TP53. Only genes with at least one drug-gene association were visualized.

### Gene set enrichment comparisons

Curated gene sets derived from Gene Ontology and the Kyoto Encyclopedia of Genes and Genomes were downloaded from the MSigDB website, version 6.1 (https://software.broadinstitute.org/gsea/msigdb/). All pairs of human genes were assessed for co-annotation by any ontology term. Aggregate enrichment of functional genes for these co-annotated gene pairs (Figure 5a) was calculated as

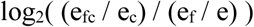

Where e_fc_ is the number of gene pairs that are both co-functional and co-annotated, e_c_ is the number of gene pairs that are co-annotated, e_f_ is the number of gene pairs that are co-functional, and e is the number of total gene pairs.

Several gene sets consist primarily of essential genes with large numbers of co-functional gene edges. Under these conditions, it is possible for a small number of genes or pathways with a large enrichment of co-functional gene edges to underlie a genome-wide enrichment, even if most pathways contain few or no co-functional genes. To estimate the number of gene sets contributing to the aggregate enrichment, we first filtered out gene sets with under 5 co-functional gene edges. These low-signal, sparsely connected gene sets are in some cases statistically enriched but would have limited use in interpreting or modeling of genome-wide networks. The remaining gene sets spanned slightly under half of GO biological process and molecular function gene sets and approximately 70% of GO cellular component and KEGG pathway sets.

Genes in the dataset were then placed into one of 100 bins based on their degree in the co-functional gene network. For each gene set, new, random gene sets were constructed such that each gene was replaced by a random gene from the same degree bin. The number of edges in the corresponding random subgraphs of the genome-wide network were calculated to serve as an empirical null distribution. If the random gene sets never reached the number of edges seen in a true gene set, a p-value of 0.5 / (# permutations) was assigned to the gene set. Significantly enriched gene sets were called at a false discovery rate of 5%. We found that over 75% of the remaining GO biological process and molecular function terms exhibited more edges than expected, and more than 90% of the remaining GO cellular component and KEGG pathways.

### Genetic interaction and protein complex comparisons

Protein-protein and genetic interaction data was downloaded from the STRING v10.5 website (detailed and action files, https://string-db.org/cgi/download.pl). Distances between gene pairs were calculated using the *distances* function from the igraph R package. The rate at which co-functional interactions were called per path length was calculated as (# co-functional interactions) divided by (# total gene pairs).

CORUM core complexes were downloaded from the CORUM website (http://mips.helmholtz-muenchen.de/corum/#download). A dataset of protein complex edges was created by enumerating all pairs of genes that were members of the same human protein complex. Across all such edges, 50.1% were called as co-functional. The expected binomial distribution per complex (Figure 5d,e) was calculated using the *dbinom* function in R, and 90% confidence intervals were calculated using *qbinom*.

To calculate Mediator gene loadings and cell line PC scores, PCA was performed on the corrected gene essentiality profiles of Mediator complex members across 342 cell lines with the *prcomp* function in R. With genes as features, the mean effect size of each gene was subtracted to estimate the covariance across genes. All loadings for the first principal component were positive, meaning all Mediator complex members covaried in the same direction, and the magnitude of the loading was taken as the relative weight for estimating the function of the entire complex.

### Community-centric analysis of the cancer cell network

Co-functional gene edges were analyzed using the igraph package in R. Distances between two genes a and b were weighted by 1 – abs(cor(gene a, gene b)) and the edge width was scaled to abs(cor(gene a, gene b)). Communities were called using the *cluster_infomap* function. The edge density of each community was calculated by constructing a subgraph from the community members and calling the *edge_density* function. In order to prevent core essential genes from influencing measures of centrality, closeness was calculated for each gene locally by calling the *closeness* function on the community subgraphs. The gene community network plot (Figure 6a) was visualized by creating a new graph of communities as nodes. Node area was made proportional to the number of genes in the community by scaling the size parameter to the square root of the number of genes. For gene communities with greater than 100 genes, the community was discarded if more than 50% of the community’s genes laid on the same chromosome. Edges were drawn between communities if the edge frequency between them surpassed the genome-wide average of 0.65% with width scaled to edge frequency.

Communities were annotated both by the number of members that were COSMIC cancer census genes (https://cancer.sanger.ac.uk/census) and enrichment in KEGG and Gene Ontology gene sets as determined by the ClusterProfiler R package. Enrichment of COSMIC census genes was determined by binomial test with probability of success equal to the total number of COSMIC census genes divided by the number of genes in the network and a false discovery rate cut-off of 10%. Enrichment of uncharacterized genes in gene clusters was determined similarly (Figure S4), where an uncharacterized gene was defined as any gene lacking a Gene Ontology biological process annotation. To understand which biological processes were sparsely connected and poorly represented (Figure S3a), communities with fewer than 8 members were analyzed using gene set enrichment.

The importance of the local closeness measure was first evaluated using the Human Protein Atlas pathology data (https://www.proteinatlas.org/about/download). For both favorable and unfavorable prognoses, genes were divided into four tiers: prognostic for 0, 1, 2, or 3+ cancer types. The Wilcoxon rank sums test was used to compare local closeness across tiers (Figure 7c). Prediction of germline How probability of loss-of-function intolerance (pLI) scores varied by closeness was also tested (http://exac.broadinstitute.org/downloads, file “fordist_cleaned_exac_r03_march16_z_pli_rec_null_data.txt”), but was weakly predictive compared to gene expression (Figure S4a).

### Gene community scores

Gene communities were processed by performing PCA on the transpose of the corrected gene essentiality profile matrix and recording the first principal component, as performed on the Mediator complex. Cell lines were scored by their first principal component score. Cell lines with large scores are interpreted as the drivers of the gene community. To examine cancer lineage and mutation status contributions to the *NF1* gene community (Figure 7d), community scores were aggregated by whether the cell lineage was glioma and whether the cell line possessed a non-silent mutation in *NF1*. To examine cancer lineage and mutation status contribution to the *KRAS* gene community (Figure 7e), community scores were aggregated by every lineage with more than 5 samples and whether the cell line possessed a missense mutation in *KRAS*. Significance of cell lineage contributions across all communities (Figure S5b) was determined by permuting cell lineage labels and calculating mean community scores per cell lineage to generate a null distribution. Cell lineages that had more extreme scores than expected were called at a 1% FDR cutoff. The gene communities with the thirty most significant associations were visualized.

### Co-functional interaction examples

The chromosome idiogram in the gene regulation example was downloaded from the Human Genome Idiogram Vector Art library (https://github.com/RCollins13/HumanIdiogramLibrary). The chromosome 17 idiogram ai file was chosen for illustration.

## Acknowledgements

We thank Michael Haney, Michael Wainburg and Michael Bassik for advice on CRISPR screen data analysis. We also thank Eilon Sharon, Emily Glassberg, Nasa Sinnott-Armstrong and other Pritchard and Greenleaf lab members for helpful discussions. This work was supported by the NIH (P50HG007735, RO1 HG008140). Evan Boyle is supported by the National Science Foundation Graduate Research Fellowship. William Greenleaf is an investigator of the Chan-Zuckerberg Biohub. Jonathan Pritchard is an investigator of the Howard Hughes Medical Institute.

## Contributions

E.A.B. conceived the project, performed analyses, and drafted the manuscript. J.K.P. and W.J.G. supervised analyses. All authors edited and revised the text.

## Supplemental figures

**Figure S1:**
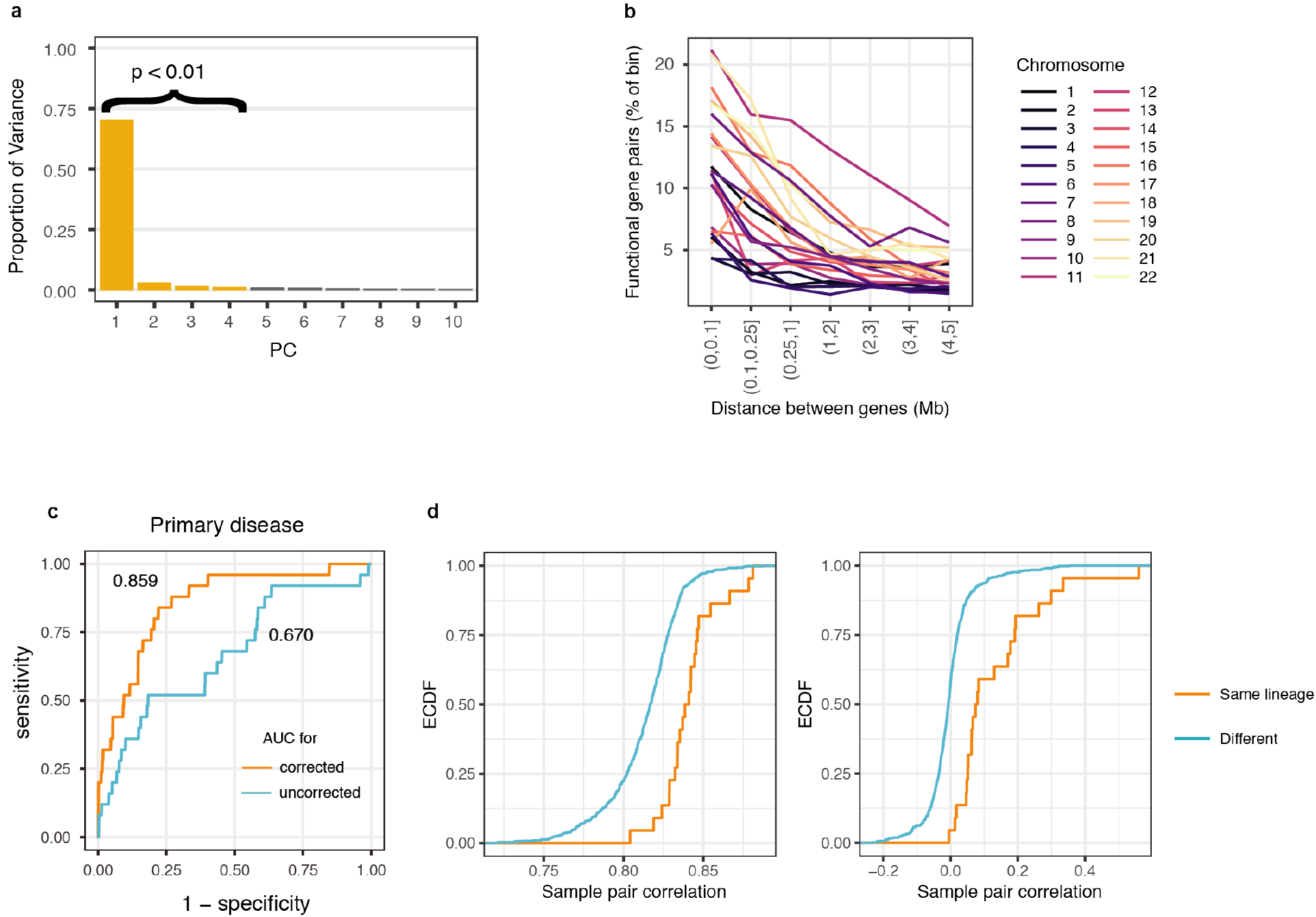
Characterization of the correction for gene essentiality profiles. **a)** PCA on olfactory receptors identifies 4 significant principal components by permutation. 68% of the variation is explained by the first principal component. **b)** The rate of calling co-functional gene pairs varies by chromosome and physical distance. **c)** Area under the receiver operating characteristic (ROC) curve for detecting two samples of the same primary disease. **d)** The empirical distributions of cell line profile pairwise correlations before and after regressing the four principal components from **a)** differ in mean.

**Figure S2:**
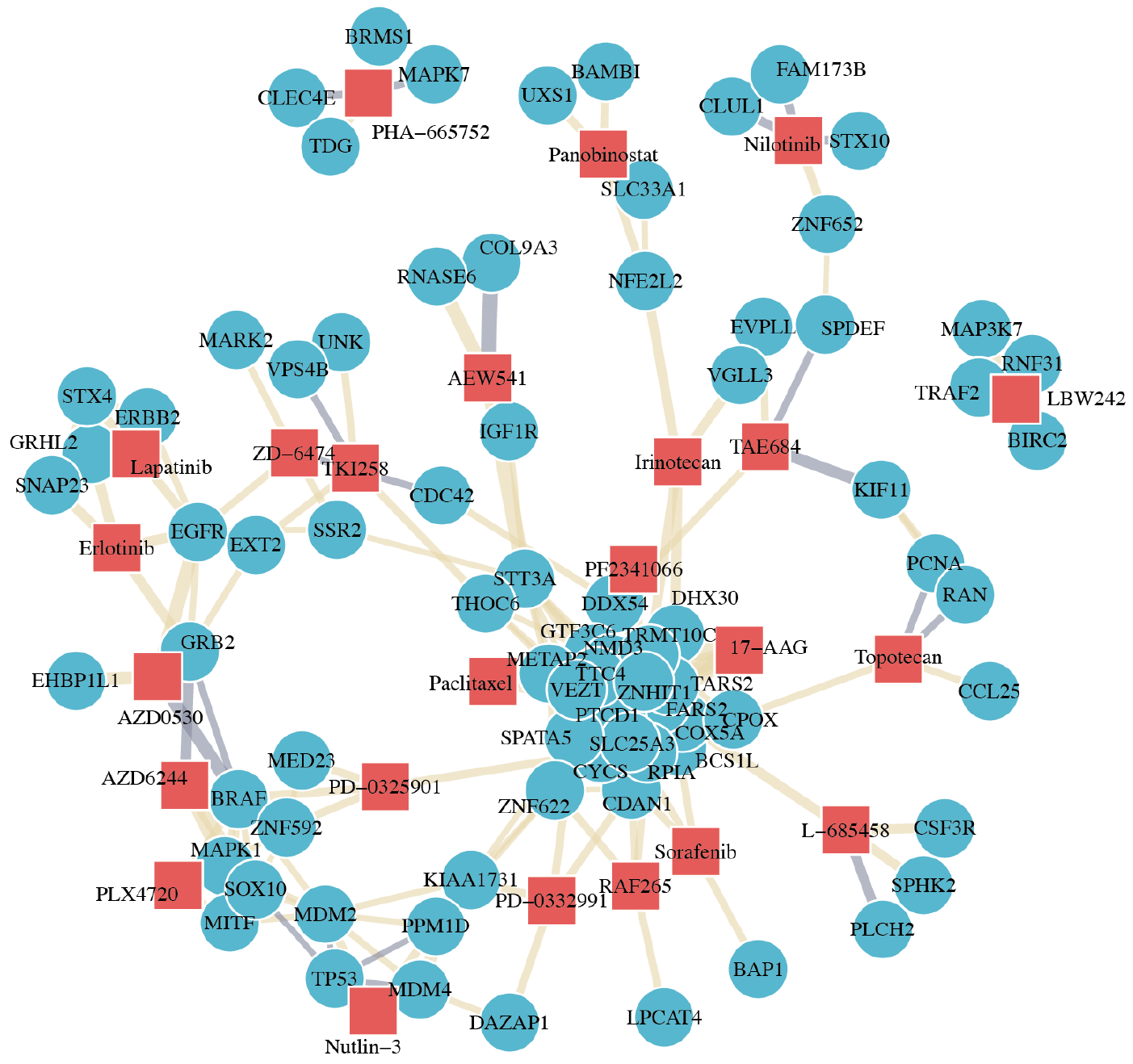
Drug-gene network. Drugs and their most correlated genes by corrected gene essentiality profiles. Correlations between gene profiles and drug profiles across cell lines creates biologically meaningful clusters for drug targets such as ErbB family members and MAPK pathways.

**Figure S3:**
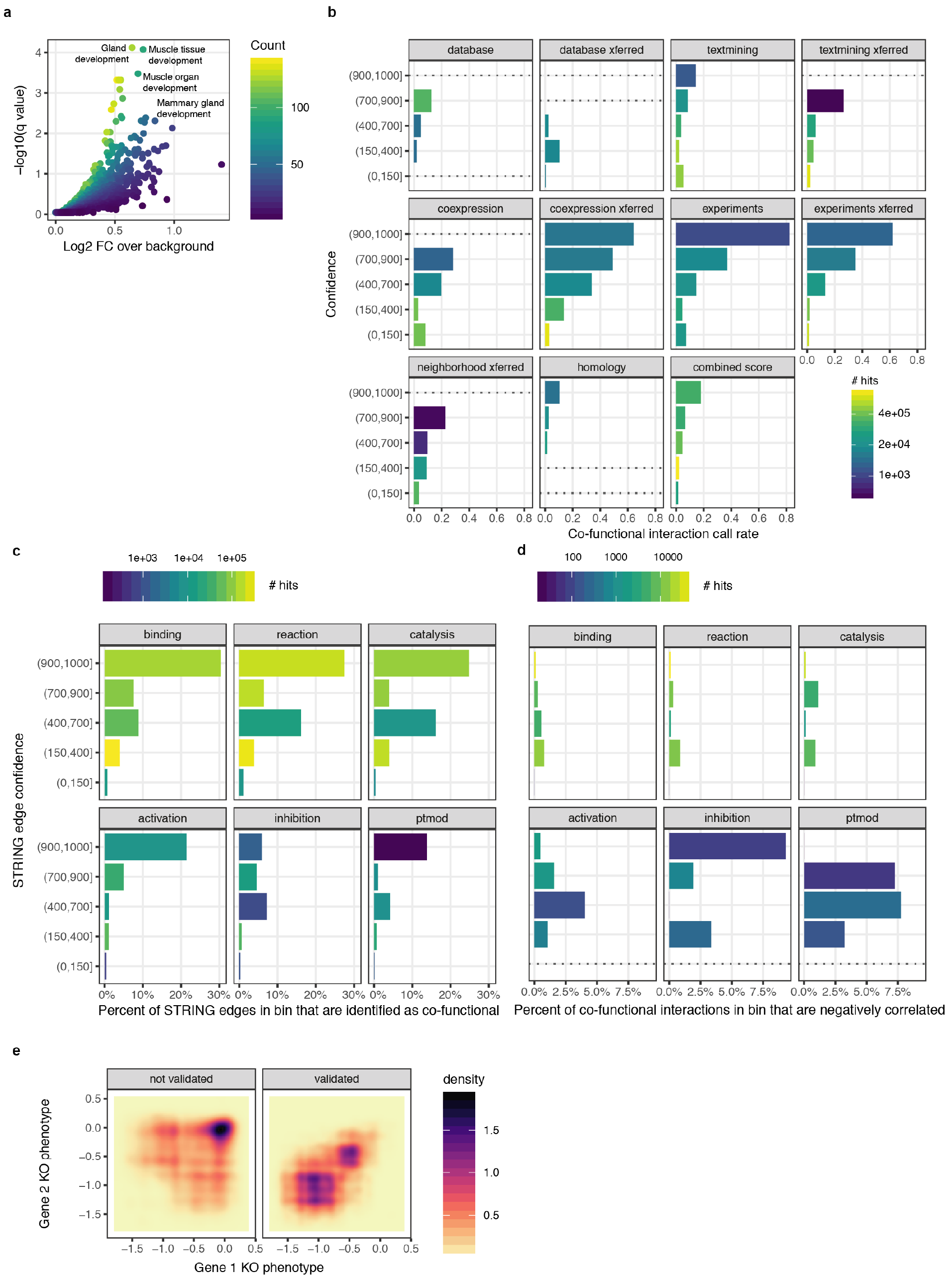
Properties of the co-functional gene network with respect to gene annotation databases. **a)** Enrichment of disconnected genes in the network across Gene Ontology Biological Process gene sets highlights terms specifying tissue differentiation and development. **b)** Co-functionality rates binned by both STRING interaction type and confidence level. “xferred” (transferred) reflects associations mapped from non-human organisms. **c)** Co-functionality rates for genetic interactions annotated with a detailed interaction type in STRING, as in **a). d)** Percent of co-functional gene edges that derived from negatively correlated corrected gene essentiality profiles, binned as in **c). e)** Density of pairs of genes participating in CORUM core complexes by mean knockout phenotype of each gene. Gene pairs that are not called as co-functional are on the left, validated gene pairs are on the right. Gene pairs with no growth deficit upon knockout are generally not called as co-functional.

**Figure S4:**
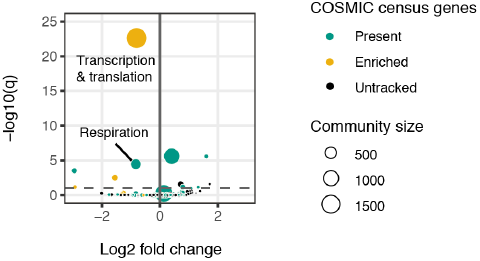
Uncharacterized genes in the cancer cell network. Volcano plot of uncharacterized gene enrichment in gene communities. Gold nodes are gene communities containing more COSMIC cancer census genes than expected by random assortment. Green nodes are gene communities containing at least one COSMIC cancer census gene.

**Figure S5:**
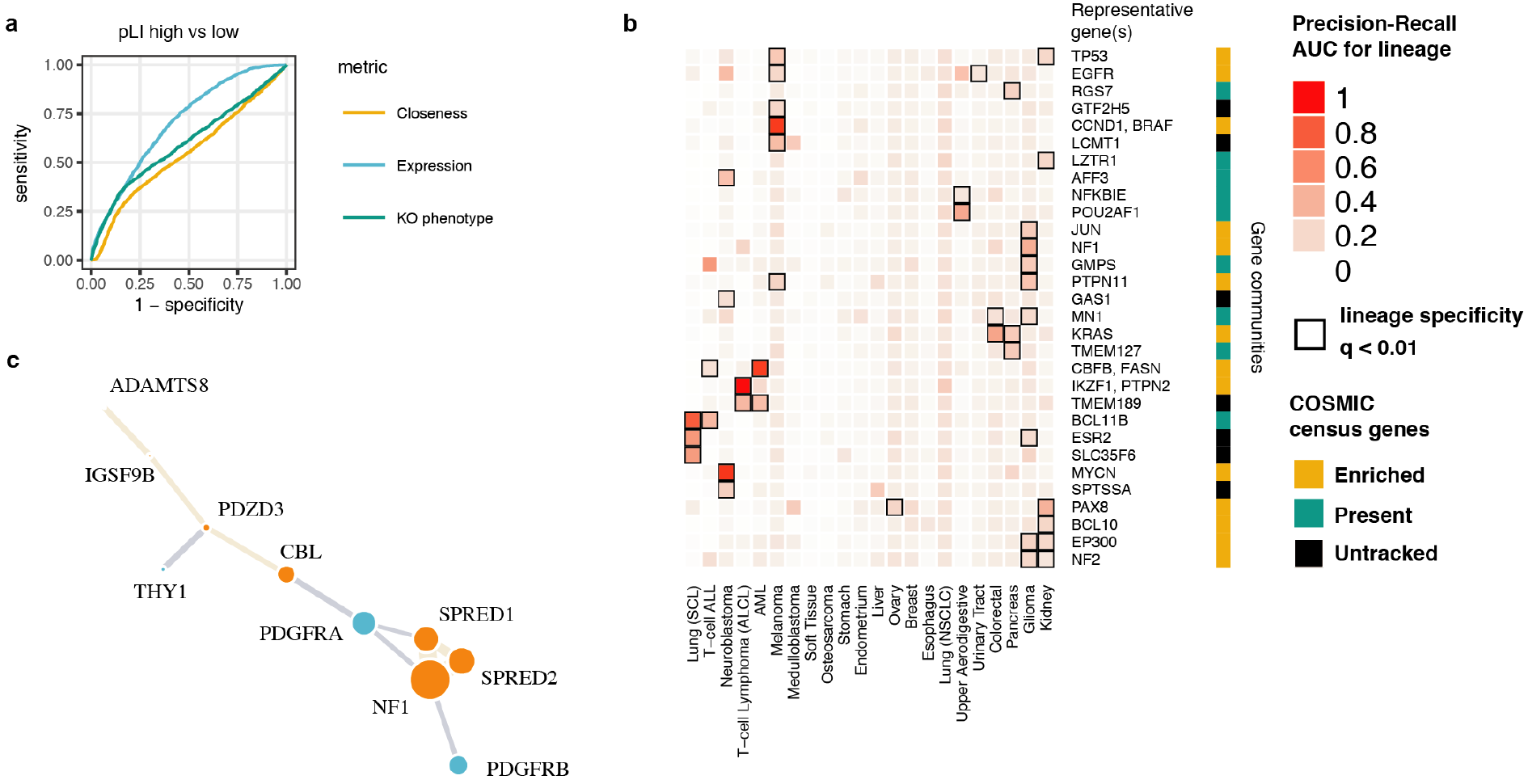
Variation in topology and cell type specificity of gene communities. **a)** ROC curve showing that loss of function-intolerant (high pLI) genes are distinguished by high expression and only weakly by growth knockout phenotype and network topology. **b)** Community scores calculated de novo from the co-functionality network often describe cell-type specific cancer regulation, especially true for gene communities enriched for COSMIC census genes (in gold). **c)** Full depiction of the NF1 gene community. Size of the gene nodes reflects loading on the first principal component amongst community members.

## References

Aguirre AJ, Meyers RM, Weir BA, Vazquez F, Zhang C-Z, Ben-David U, Cook A, Ha G, Harrington WF, Doshi MB, Kost-Alimova M, Gill S, Xu H, Ali LD, Jiang G, Pantel S, Lee Y, Goodale A, Cherniack AD, Oh C, et al (2016) Genomic Copy Number Dictates a Gene-Independent Cell Response to CRISPR/Cas9 Targeting. Cancer Discov 6: 914–929

Ashburner M, Ball CA, Blake JA, Botstein D, Butler H, Cherry JM, Davis AP, Dolinski K, Dwight SS, Eppig JT, Harris MA, Hill DP, Issel-Tarver L, Kasarskis A, Lewis S, Matese JC, Richardson JE, Ringwald M, Rubin GM & Sherlock G (2000) Gene ontology: tool for the unification of biology. The Gene Ontology Consortium. Nat Genet 25: 25–29

Barabási A-L & Oltvai ZN (2004) Network biology: understanding the cell’s functional organization. Nat. Rev. Genet. 5: 101–113

Barabási A-L, Gulbahce N & Loscalzo J (2011) Network medicine: a network-based approach to human disease. Nat. Rev. Genet. 12: 56–68

Barretina J, Caponigro G, Stransky N, Venkatesan K, Margolin AA, Kim S, Wilson CJ, Lehár J, Kryukov GV, Sonkin D, Reddy A, Liu M, Murray L, Berger MF, Monahan JE, Morais P, Meltzer J, Korejwa A, Jané-Valbuena J, Mapa FA, et al (2012) The Cancer Cell Line Encyclopedia enables predictive modelling of anticancer drug sensitivity. Nature 483: 603–607

Bassik MC, Kampmann M, Lebbink RJ, Wang S, Hein MY, Poser I, Weibezahn J, Horlbeck MA, Chen S, Mann M, Hyman AA, LeProust EM, McManus MT & Weissman JS (2013) A systematic mammalian genetic interaction map reveals pathways underlying ricin susceptibility. Cell 152: 909–922

Bovée JVMG (2008) Multiple osteochondromas. Orphanet J Rare Dis 3: 3

Breeze CE, Paul DS, van Dongen J, Butcher LM, Ambrose JC, Barrett JE, Lowe R, Rakyan VK, Iotchkova V, Frontini M, Downes K, Ouwehand WH, Laperle J, Jacques P-É, Bourque G, Bergmann AK, Siebert R, Vellenga E, Saeed S, Matarese F, et al (2016) eFORGE: A Tool for Identifying Cell Type-Specific Signal in Epigenomic Data. Cell Rep 17: 2137–2150

Costanzo M, Baryshnikova A, Bellay J, Kim Y, Spear ED, Sevier CS, Ding H, Koh JLY, Toufighi K, Mostafavi S, Prinz J, St Onge RP, VanderSluis B, Makhnevych T, Vizeacoumar FJ, Alizadeh S, Bahr S, Brost RL, Chen Y, Cokol M, et al (2010) The genetic landscape of a cell. Science 327: 425–431

Costanzo M, VanderSluis B, Koch EN, Baryshnikova A, Pons C, Tan G, Wang W, Usaj M, Hanchard J, Lee SD, Pelechano V, Styles EB, Billmann M, van Leeuwen J, Van Dyk N, Lin Z-Y, Kuzmin E, Nelson J, Piotrowski JS, Srikumar T, et al (2016) A global genetic interaction network maps a wiring diagram of cellular function. Science 353:

Deshpande R, Nelson J, Simpkins SW, Costanzo M, Piotrowski JS, Li SC, Boone C & Myers CL Efficient strategies for screening large-scale genetic interaction networks.

Doench JG, Fusi N, Sullender M, Hegde M, Vaimberg EW, Donovan KF, Smith I, Tothova Z, Wilen C, Orchard R, Virgin HW, Listgarten J & Root DE (2016) Optimized sgRNA design to maximize activity and minimize off-target effects of CRISPR-Cas9. Nat. Biotechnol. 34: 184–191

Fontana L, Partridge L & Longo VD (2010) Extending Healthy Life Span--From Yeast to Humans. Science 328: 321–326

Forbes SA, Beare D, Boutselakis H, Bamford S, Bindal N, Tate J, Cole CG, Ward S, Dawson E, Ponting L, Stefancsik R, Harsha B, Kok CY, Jia M, Jubb H, Sondka Z, Thompson S, De T & Campbell PJ (2017) COSMIC: somatic cancer genetics at high-resolution. Nucleic Acids Research 45: D777–D783

Fortunato S (2010) Community detection in graphs. Physics Reports 486: 75–174

Gavory G, O’Dowd CR, Helm MD, Flasz J, Arkoudis E, Dossang A, Hughes C, Cassidy E, McClelland K, Odrzywol E, Page N, Barker O, Miel H & Harrison T (2018) Discovery and characterization of highly potent and selective allosteric USP7 inhibitors. Nat. Chem. Biol. 14: 118–125

Gilbert LA, Horlbeck MA, Adamson B, Villalta JE, Chen Y, Whitehead EH, Guimaraes C, Panning B, Ploegh HL, Bassik MC, Qi LS, Kampmann M & Weissman JS (2014) Genome-Scale CRISPR-Mediated Control of Gene Repression and Activation. Cell 159: 647–661

Gilmore TD (2006) Introduction to NF-κB: players, pathways, perspectives. Oncogene 25: 6680–6684

GTEx Consortium, Laboratory, Data Analysis &Coordinating Center (LDACC)—Analysis Working Group, Statistical Methods groups—Analysis Working Group, Enhancing GTEx (eGTEx) groups, NIH Common Fund, NIH/NCI, NIH/NHGRI, NIH/NIMH, NIH/NIDA, Biospecimen Collection Source Site—NDRI, Biospecimen Collection Source Site—RPCI, Biospecimen Core Resource— VARI, Brain Bank Repository—University of Miami Brain Endowment Bank, Leidos Biomedical— Project Management, ELSI Study, Genome Browser Data Integration &Visualization—EBI, Genome Browser Data Integration &Visualization—UCSC Genomics Institute, University of California Santa Cruz, Lead analysts:, Laboratory, Data Analysis &Coordinating Center (LDACC):, NIH program management:, et al (2017) Genetic effects on gene expression across human tissues. Nature 550: 204–213

Gutmann DH, Ferner RE, Listernick RH, Korf BR, Wolters PL & Johnson KJ (2017) Neurofibromatosis type 1. Nat Rev Dis Primers 3: 17004

Han K, Jeng EE, Hess GT, Morgens DW, Li A & Bassik MC (2017) Synergistic drug combinations for cancer identified in a CRISPR screen for pairwise genetic interactions. Nat. Biotechnol. 35: 463–474

Hart T, Brown KR, Sircoulomb F, Rottapel R & Moffat J (2014) Measuring error rates in genomic perturbation screens: gold standards for human functional genomics. Mol. Syst. Biol. 10: 733

Hart T, Chandrashekhar M, Aregger M, Steinhart Z, Brown KR, MacLeod G, Mis M, Zimmermann M, Fradet-Turcotte A, Sun S, Mero P, Dirks P, Sidhu S, Roth FP, Rissland OS, Durocher D, Angers S & Moffat J (2015) High-Resolution CRISPR Screens Reveal Fitness Genes and Genotype-Specific Cancer Liabilities. Cell 163: 1515–1526

Javierre BM, Burren OS, Wilder SP, Kreuzhuber R, Hill SM, Sewitz S, Cairns J, Wingett SW, Várnai C, Thiecke MJ, Burden F, Farrow S, Cutler AJ, Rehnström K, Downes K, Grassi L, Kostadima M, Freire-Pritchett P, Wang F, BLUEPRINT Consortium, et al (2016) Lineage-Specific Genome Architecture Links Enhancers and Non-coding Disease Variants to Target Gene Promoters. Cell 167: 1369–1384.e19

Kampmann M, Horlbeck MA, Chen Y, Tsai JC, Bassik MC, Gilbert LA, Villalta JE, Kwon SC, Chang H, Kim VN & Weissman JS (2015) Next-generation libraries for robust RNA interference-based genome-wide screens. Proc. Natl. Acad. Sci. U.S.A. 112: E3384–91

Kim E, Dede M, Lenoir WF, Wang G, Srinivasan S, Colic M & Hart T (2018) Hierarchical organization of the human cell from a cancer coessentiality network.

Lek M, Karczewski KJ, Minikel EV, Samocha KE, Banks E, Fennell T, O’Donnell-Luria AH, Ware JS, Hill AJ, Cummings BB, Tukiainen T, Birnbaum DP, Kosmicki JA, Duncan LE, Estrada K, Zhao F, Zou J, Pierce-Hoffman E, Berghout J, Cooper DN, et al (2016) Analysis of protein-coding genetic variation in 60,706 humans. Nature 536: 285–291

Lindsey S & Langhans SA (2015) Epidermal growth factor signaling in transformed cells. Int Rev Cell Mol Biol 314: 1–41

Medina PJ & Goodin S (2008) Lapatinib: a dual inhibitor of human epidermal growth factor receptor tyrosine kinases. Clin Ther 30: 1426–1447

Meyers RM, Bryan JG, McFarland JM, Weir BA, Sizemore AE, Xu H, Dharia NV, Montgomery PG, Cowley GS, Pantel S, Goodale A, Lee Y, Ali LD, Jiang G, Lubonja R, Harrington WF, Strickland M, Wu T, Hawes DC, Zhivich VA, et al (2017) Computational correction of copy number effect improves specificity of CRISPR-Cas9 essentiality screens in cancer cells. Nat Genet 49: 1779–1784

Morgens DW, Wainberg M, Boyle EA, Ursu O, Araya CL, Tsui CK, Haney MS, Hess GT, Han K, Jeng EE, Li A, Snyder MP, Greenleaf WJ, Kundaje A & Bassik MC (2017) Genome-scale measurement of off-target activity using Cas9 toxicity in high-throughput screens. Nat Commun 8: 15178

Mumbach MR, Satpathy AT, Boyle EA, Dai C, Gowen BG, Cho SW, Nguyen ML, Rubin AJ, Granja JM, Kazane KR, Wei Y, Nguyen T, Greenside PG, Corces MR, Tycko J, Simeonov DR, Suliman N, Li R, Xu J, Flynn RA, et al (2017) Enhancer connectome in primary human cells identifies target genes of disease-associated DNA elements. Nat Genet 49: 1602–1612

Ogasawara S, Kiyota Y, Chuman Y, Kowata A, Yoshimura F, Tanino K, Kamada R & Sakaguchi K (2015) Novel inhibitors targeting PPM1D phosphatase potently suppress cancer cell proliferation. Bioorg. Med. Chem. 23: 6246–6249

Ornitz DM & Itoh N (2015) The Fibroblast Growth Factor signaling pathway. Wiley Interdiscip Rev Dev Biol 4: 215–266

Pan J, Meyers RM, Michel BC, Mashtalir N, Sizemore AE, Wells JN, Cassel SH, Vazquez F, Weir BA, Hahn WC, Marsh JA, Tsherniak A & Kadoch C (2018) Interrogation of Mammalian Protein Complex Structure, Function, and Membership Using Genome-Scale Fitness Screens. Cell Syst 6: 555–568.e7

Park RJ, Wang T, Koundakjian D, Hultquist JF, Lamothe-Molina P, Monel B, Schumann K, Yu H, Krupzcak KM, Garcia-Beltran W, Piechocka-Trocha A, Krogan NJ, Marson A, Sabatini DM, Lander ES, Hacohen N & Walker BD (2017) A genome-wide CRISPR screen identifies a restricted set of HIV host dependency factors. Nat Genet 49: 193–203

Pickrell JK, Marioni JC, Pai AA, Degner JF, Engelhardt BE, Nkadori E, Veyrieras J-B, Stephens M, Gilad Y & Pritchard JK (2010) Understanding mechanisms underlying human gene expression variation with RNA sequencing. Nature 464: 768–772

Pommier Y (2006) Topoisomerase I inhibitors: camptothecins and beyond. Nat. Rev. Cancer 6: 789–802

Rhee SY, Wood V, Dolinski K & Draghici S (2008) Use and misuse of the gene ontology annotations. Nat. Rev. Genet. 9: 509–515

Roadmap Epigenomics Consortium, Kundaje A, Meuleman W, Ernst J, Bilenky M, Yen A, Heravi-Moussavi A, Kheradpour P, Zhang Z, Wang J, Ziller MJ, Amin V, Whitaker JW, Schultz MD, Ward LD, Sarkar A, Quon G, Sandstrom RS, Eaton ML, Wu Y-C, et al (2015) Integrative analysis of 111 reference human epigenomes. Nature 518: 317–330

Rolland T, Taşan M, Charloteaux B, Pevzner SJ, Zhong Q, Sahni N, Yi S, Lemmens I, Fontanillo C, Mosca R, Kamburov A, Ghiassian SD, Yang X, Ghamsari L, Balcha D, Begg BE, Braun P, Brehme M, Broly MP, Carvunis A-R, et al (2014) A proteome-scale map of the human interactome network. Cell 159: 1212–1226

Rosenbluh J, Xu H, Harrington W, Gill S, Wang X, Vazquez F, Root DE, Tsherniak A & Hahn WC (2017) Complementary information derived from CRISPR Cas9 mediated gene deletion and suppression. Nat Commun 8: 15403

Ruepp A, Waegele B, Lechner M, Brauner B, Dunger-Kaltenbach I, Fobo G, Frishman G, Montrone C & Mewes H-W (2010) CORUM: the comprehensive resource of mammalian protein complexes--2009. Nucleic Acids Research 38: D497–501

Shalem O, Sanjana NE, Hartenian E, Shi X, Scott DA, Mikkelsen TS, Heckl D, Ebert BL, Root DE, Doench JG & Zhang F (2014) Genome-scale CRISPR-Cas9 knockout screening in human cells. Science 343: 84–87

Shen JP, Zhao D, Sasik R, Luebeck J, Birmingham A, Bojorquez-Gomez A, Licon K, Klepper K, Pekin D, Beckett AN, Sanchez KS, Thomas A, Kuo C-C, Du D, Roguev A, Lewis NE, Chang AN, Kreisberg JF, Krogan N, Qi L, et al (2017) Combinatorial CRISPR-Cas9 screens for de novo mapping of genetic interactions. Nat Methods 14: 573–576

Stumpf MPH, Thorne T, de Silva E, Stewart R, An HJ, Lappe M & Wiuf C (2008) Estimating the size of the human interactome. Proc. Natl. Acad. Sci. U.S.A. 105: 6959–6964

Subramanian A, Tamayo P, Mootha VK, Mukherjee S, Ebert BL, Gillette MA, Paulovich A, Pomeroy SL, Golub TR, Lander ES & Mesirov JP (2005) Gene set enrichment analysis: a knowledge-based approach for interpreting genome-wide expression profiles. PNAS 102: 15545–15550

Szklarczyk D, Franceschini A, Wyder S, Forslund K, Heller D, Huerta-Cepas J, Simonovic M, Roth A, Santos A, Tsafou KP, Kuhn M, Bork P, Jensen LJ & Mering von C (2015) STRING v10: protein-protein interaction networks, integrated over the tree of life. Nucleic Acids Research 43: D447–52

The Gene Ontology Consortium (2017) Expansion of the Gene Ontology knowledgebase and resources. Nucleic Acids Research 45: D331–D338

Timmons JA, Szkop KJ & Gallagher IJ (2015) Multiple sources of bias confound functional enrichment analysis of global-omics data. Genome Biol. 16: 186

Tong AHY, Lesage G, Bader GD, Ding H, Xu H, Xin X, Young J, Berriz GF, Brost RL, Chang M, Chen Y, Cheng X, Chua G, Friesen H, Goldberg DS, Haynes J, Humphries C, He G, Hussein S, Ke L, et al (2004) Global mapping of the yeast genetic interaction network. Science 303: 808–813

Uhlen M, Zhang C, Lee S, Sjöstedt E, Fagerberg L, Bidkhori G, Benfeitas R, Arif M, Liu Z, Edfors F, Sanli K, Feilitzen von K, Oksvold P, Lundberg E, Hober S, Nilsson P, Mattsson J, Schwenk JM, Brunnström H, Glimelius B, et al (2017) A pathology atlas of the human cancer transcriptome. Science 357:

Venkatesan K, Rual J-F, Vazquez A, Stelzl U, Lemmens I, Hirozane-Kishikawa T, Hao T, Zenkner M, Xin X, Goh K-I, Yildirim MA, Simonis N, Heinzmann K, Gebreab F, Sahalie JM, Cevik S, Simon C, de Smet A-S, Dann E, Smolyar A, et al (2009) An empirical framework for binary interactome mapping. Nat Methods 6: 83–90

Wang T, Birsoy K, Hughes NW, Krupczak KM, Post Y, Wei JJ, Lander ES & Sabatini DM (2015) Identification and characterization of essential genes in the human genome. Science 350: 1096–1101

Wang T, Wei JJ, Sabatini DM & Lander ES (2014) Genetic screens in human cells using the CRISPR-Cas9 system. Science 343: 80–84

Wang T, Yu H, Hughes NW, Liu B, Kendirli A, Klein K, Chen WW, Lander ES & Sabatini DM (2017) Gene Essentiality Profiling Reveals Gene Networks and Synthetic Lethal Interactions with Oncogenic Ras. Cell 168: 890–903.e15

Wong ASL, Choi GCG, Cui CH, Pregernig G, Milani P, Adam M, Perli SD, Kazer SW, Gaillard A, Hermann M, Shalek AK, Fraenkel E & Lu TK (2016) Multiplexed barcoded CRISPR-Cas9 screening enabled by CombiGEM. Proc. Natl. Acad. Sci. U.S.A. 113: 2544–2549

Wu A, Xiao T, Fei T, Liu SX & Li W Reducing False Positives in CRISPR/Cas9 Screens from Copy Number Variations.

Yin J-W & Wang G (2014) The Mediator complex: a master coordinator of transcription and cell lineage development. Development 141: 977–987

Zhang Y & Lu H (2009) Signaling to p53: ribosomal proteins find their way. Cancer Cell 16: 369–377

